# Sleep enhances spatial schema memory formation in humans

**DOI:** 10.64898/2026.06.16.732347

**Authors:** Lisa Bastian, Hannah Hamann, Tim Näher, Karsten Rauss, Jan Born

## Abstract

Schema memory, a generalized representation formed across episodes sharing regularities, is thought to arise through sleep-dependent systems consolidation. Yet, direct evidence in humans remains sparse. Here, sixty young adults navigated a virtual-reality arena and learned a spatial distribution of object-category ratios across five sessions (toys vs. household items, hidden in boxes at different locations). Participants then either slept or were sleep-deprived for a full night, followed by two recovery nights before memory testing, or they were tested after a short 30-min delay spent awake. Only after a three-day delay did spatial memory of old box locations predict spatial interpolation to new box locations, indicating time-dependent schema expression. Critically, sleep distinctly enhanced spatial integration beyond the effect of time. This benefit was predicted by frontal cortical slow oscillation-spindle coupling during the first post-encoding night, thus linking sleep oscillations to the transformation of episodic spatial memories into integrated schema representations.

## Introduction

Schema memory refers to generalized knowledge structures that capture regularities across multiple episodes. These structures serve as flexible templates for interpreting and acting upon novel situations^1–4^. For instance, when visiting a new town, we effortlessly locate the town hall without being told where it is by drawing on generalized knowledge about how cities tend to be arranged. Unlike episodic memory, which preserves the specific content and spatiotemporal context of individual events, schemas distill structural invariants shared across experiences into representations that transcend any single episode. Schemas should also be distinguished from gist information, i.e., the central elements of a single episode stripped of peripheral detail^5,6^. A schema integrates the gist of multiple episodes to represent a regularity that generalizes across episodes^2,3^.

In humans, de novo formation of schemas has predominantly been studied using category learning and temporal dependencies; for reviews see^2,7,8^. Some studies have used computerized object-space associations^9,10^ but true spatial schema formation, that is, generalized representations of recurring spatial regularities formed through navigational experience^4^, has rarely been investigated in humans^11–13^. This is a surprising gap, given that influential animal work on schemas was conducted using real spatial paradigms that involve navigation^14–17^. Studying spatial schema formation in humans requires paradigms that control for pre-existing knowledge while engaging the allocentric spatial processing central to much of the research in animal models. Virtual reality (VR) offers a powerful solution, enabling participants to actively navigate large-scale, controlled environments ^18–20^ in which a spatial rule can be embedded across multiple learning episodes without any prior familiarity.

Schema formation is considered to be a gradual process that evolves over time^21–24^. It remains an open question whether sleep plays an active role in schema formation. Active systems consolidation theory proposes a conclusive mechanism: During non-rapid eye-movement (NREM) sleep, newly encoded hippocampal-dependent representations are repeatedly reactivated and strengthened in neocortical long-term stores^25–28^. The hippocampal-neocortical dialogue underlying this representational change is orchestrated by the temporal coupling of neocortical slow oscillations (SOs), thalamocortical sleep spindles, and hippocampal sharp-wave ripples^25,29^ with the precise nesting of spindles within the SO up-state (i.e. SO-spindle coupling) opening windows of enhanced neuroplasticity in neocortical circuits that mediate the long-term storage of memories^30–32^.

Importantly, beyond strengthening individual memory traces, this process is thought to qualitatively transform representations: repeated reactivation of overlapping memories during NREM sleep gradually strengthens shared features while weakening idiosyncratic details, which may foster the formation of schema-like representations^27,33,34^. Consistent with this idea, sleep facilitates the detection of hidden spatial regularities^35,36^ and the integration of new information with prior knowledge^37,38^. Moreover, evidence in rodents and humans suggests that memory hippocampal reactivations during NREM sleep in particular synchronize with medial prefrontal cortical (mPFC) activity, which is thought to foster the formation of schema-like neocortical representation^15,39–43^, pointing to a particular role of the hippocampal dialogue with mPFC regions in mediating sleep-dependent schema memory formation.

Here, we used an immersive VR paradigm in which participants navigated a hexagonal arena and learned, by navigating to boxes hiding either a toy or a household item, a spatial distribution of object-category ratios across five learning sessions. After a retention interval comprising either a short 30-min delay, a full night of sleep, or a night of sleep deprivation, both followed by two recovery nights, participants recalled learned information and, in addition, inferred category ratios at novel, previously unseen box locations. This test at unseen box locations served as a spatial interpolation task to test schema memory. We used graph-theoretic analysis to quantify schema memory as the structural correspondence between participants’ response graphs and the true spatial distribution, capturing the integrated spatial representation. We tested three hypotheses. First, the passage of time is necessary for episodic memories to become usable for spatial interpolation, schema-like integration is evident only after a multi-day delay, but not after the short 30-min delay. Second, sleep enhances spatial schema memory beyond the effect of time: Participants who slept after encoding show superior local spatial integration compared with sleep-deprived participants, despite identical delays. Third, SO-spindle coupling specifically over frontal cortical regions during the first post-encoding night predicts schema memory, implicating a hippocampal-prefrontal cortical dialogue during sleep as a mechanism underlying spatial schema formation.

## Results

60 healthy young adults (mean = 23.5 years, 50% female) participated in the study. Upon introduction to the VR equipment, participants completed a 15-min guided familiarization phase in the hexagonal arena in VR (**Fig. 1a**), immediately followed by the schema build-up phase comprising five learning sessions spaced by 15-min breaks, each including an immediate retrieval (**Fig. 1d**).

**Fig. 1.**
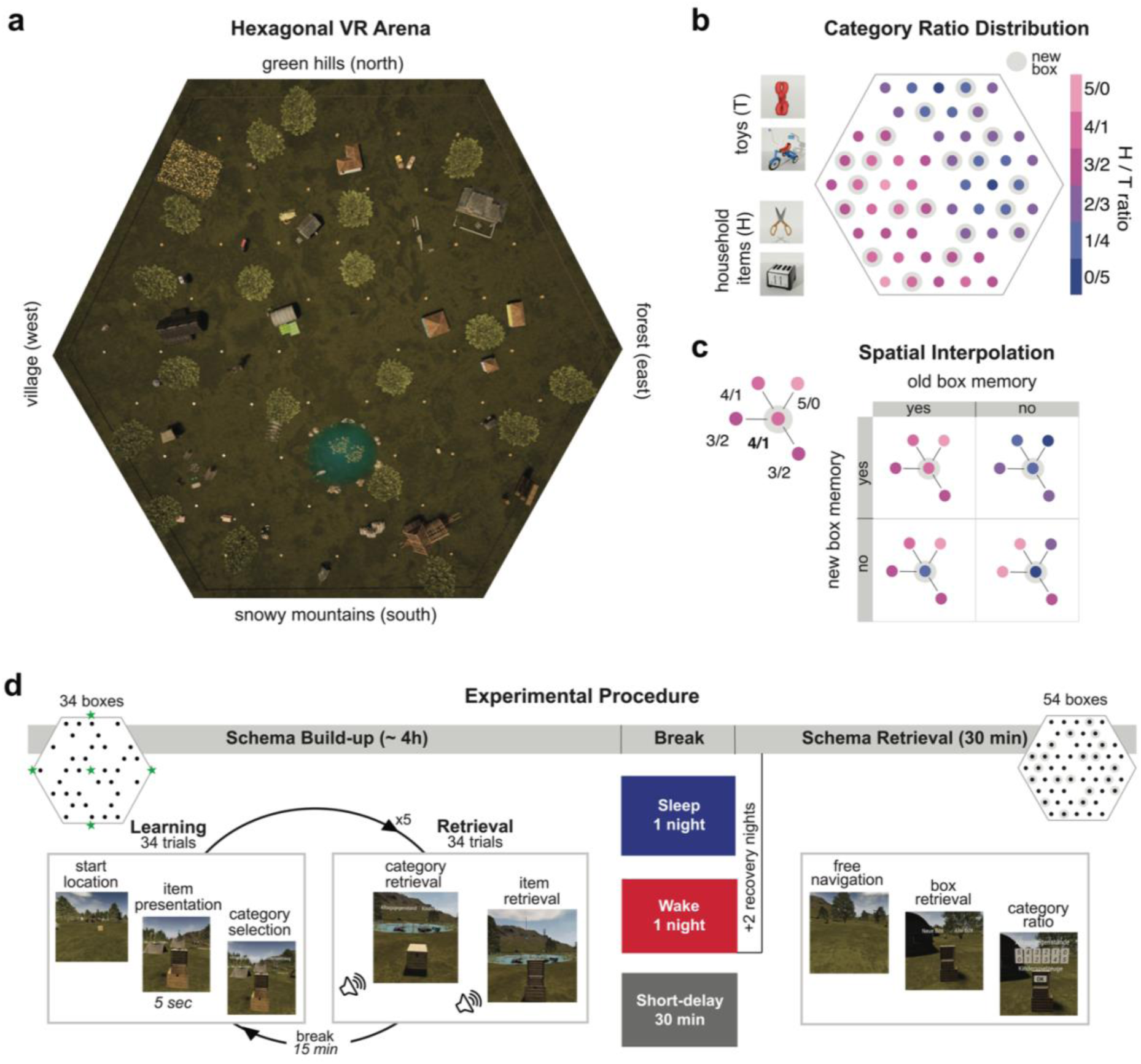
Experimental paradigm and spatial schema design. **a**, Bird’s-eye view of the hexagonal virtual-reality (VR) arena containing distal spatial cues (not shown - green hills to the north, forest to the east, snowy mountains to the south, village to the west), proximal landmarks (buildings, trees, a campground, a field, cars, a lake, a windmill and a sawmill), and 54 boxes arranged in a hexagonal pattern. **b**, Spatial category ratio distribution across the arena. All boxes (indicated by dots) looked identical. Each of the 34 old boxes contained five objects drawn from two categories (toys and household items) in varying ratios, creating a continuous spatial gradient of household-to-toy ratios (H/T ratio, color scale for dots, 5/0 to 0/5). Grey shading indicates the 20 new box locations added for the schema retrieval session. Example objects from each category are shown on the left. **c**, Spatial interpolation principle. Top left: each new box was surrounded by equidistant old-box neighbors whose category ratios determined the expected ratio at the new location (here, 4/1, computed from the rounded average of surrounding ratios). Right: the correct interpolation to new box locations (schema memory) and the correct recall of old boxes (i.e., episodic memory of build-up experience) represent orthogonal performance dimensions; participants could succeed on both, one or none of the two. **d**, Experimental procedure. During schema build-up (∼4 h), participants completed five learning sessions, each comprising 34 navigation trials (navigate to a box, view the object for 5 s, report the category) followed by 34 retrieval trials (retrieve the category and identify the correct object, followed by auditory response feedback), separated by 15-min breaks. Green stars on the arena outline indicate the different start locations used across the 5 learning sessions to promote allocentric encoding. After schema build-up, participants were assigned to one of three conditions: sleep with polysomnography (Sleep; blue), overnight sleep deprivation (Wake; red), or a short 30-min delay spent awake (Short-delay; grey). The Sleep and Wake conditions included two additional recovery nights before schema retrieval three days later; the Short-delay condition proceeded to schema retrieval after 30 min. At schema retrieval (∼30 min), participants were spawned at the arena center and freely navigated to all 54 boxes (34 old + 20 new), reported whether each box was old or new, and indicated the category ratio for each location.

During schema build-up, 34 boxes were placed in the VR environment. For each learning session, the trial start location differed to facilitate allocentric encoding of the environment. In each trial, the participant navigated freely to one of the 34 boxes, each of which contained either a household item or a toy (**Fig. 1b**). After arriving in front of a box, the hidden object was presented for 5 s and the participant subsequently responded to which category of the object belonged to (**Video S1**). The 34 navigation trials of an episode were followed by 34 trials of an immediate retrieval test. On each retrieval trial, the participant was spawned in front of one box and was asked to recall the category of the hidden object in the box (household item vs. toy). The participant was then presented with two items, one of which had been in this box during learning, and their task was to select the correct one (**Video S2**). Auditory feedback was given after each response to help the participant remember the correct category and item for each box location.

Following all five learning sessions, the participants were assigned to one of three conditions: 1) a full night of sleep in the lab with polysomnography (Sleep; *N* = 20), 2) a night of sleep deprivation in the lab (Wake; *N* = 20), a short 30-min delay followed by immediate testing (Short-delay; *N* = 20). Participants in the Sleep and Wake conditions returned to the lab three days after schema-build-up following two recovery nights (the first of which was in the sleep lab for the Wake group).

At schema retrieval, participants encountered not only the 34 old boxes from schema build-up but also 20 additional new boxes at new locations. Old-box memory served to test the spatial aspect of episodic memory formed during schema build-up, while new-box memory was used to probe spatial schema formation. Participants were spawned in the center of the arena and then freely navigated to each of the 54 boxes. For each box, they first indicated whether it was an old or new box location. For old locations, the participant then indicated the category ratio of toys to household items drawing on their episodic memory formed during schema build-up. For new boxes, they had to infer the category ratios based on the box location in the arena (**Video S3**). The correct category ratio for each new box was determined by the average ratios of its directly surrounding old boxes (**see Fig. 1c**). The ability to successfully interpolate in space indicated formation of a spatial schema, i.e., the integrated representation of the spatial category-ratio distribution.

### Memory for old box locations predicts performance for new box locations after a three-day delay

During schema build-up, category recall was not different across the three retention conditions (Sleep, Wake and Short-delay; **Fig. 2a**, *F*_2,569_ = 0.35, *P* = .703) and object categories (Household items and Toys; **Fig. S1a,** *F*_1,569_ = 2.46, *P* = .117). Participants improved in category recall across sessions (**Fig. 2a**; *F*_4,569_ = 2.52, *P* = .040, η² = 0.02), specifically in the 4^th^ session compared to the first (follow-up *t*_569_ = 2.79, *P* = .043) and marginally in the 5^th^ learning session compared to the first (follow-up *t*_569_ = 2.58, *P* = .074). This improvement is consistent with participants developing a more effective task strategy over the course of the schema build-up phase. We found no gender difference in performance (*F*_1,569_ < 0.01, *P* = .989). Similarly, item recognition was comparable across retention conditions (**Fig. S1b**, *F*_2,284_ = 0.33, *P* = .717) and gender (*F*_1,56_ = 1.29, *P* = .167). Participants recognized fewer items in the first session compared with sessions two (*t*_284_ = -4.45, *P* < .001), three (*t*_284_ = -4.45, *P* < .001), four (*t*_284_ = -6.37, *P* < .001) and five (*t*_284_ = -4.50, *P* < .001). Sustained attention, as assessed with the psychomotor vigilance test ^44^, did not differ between the retention conditions across build-up sessions and schema retrieval (**Fig. S2b**, Condition x Session: *F*_12,398_ = 0.43, *P* = .950). Control measures including self-reported sleepiness, mood, and motion sickness also indicated that, overall, experienced mental and physiological states were well-matched across conditions at schema build-up (see **Fig. S2** and Supplementary Results for details).

**Fig. 2.**
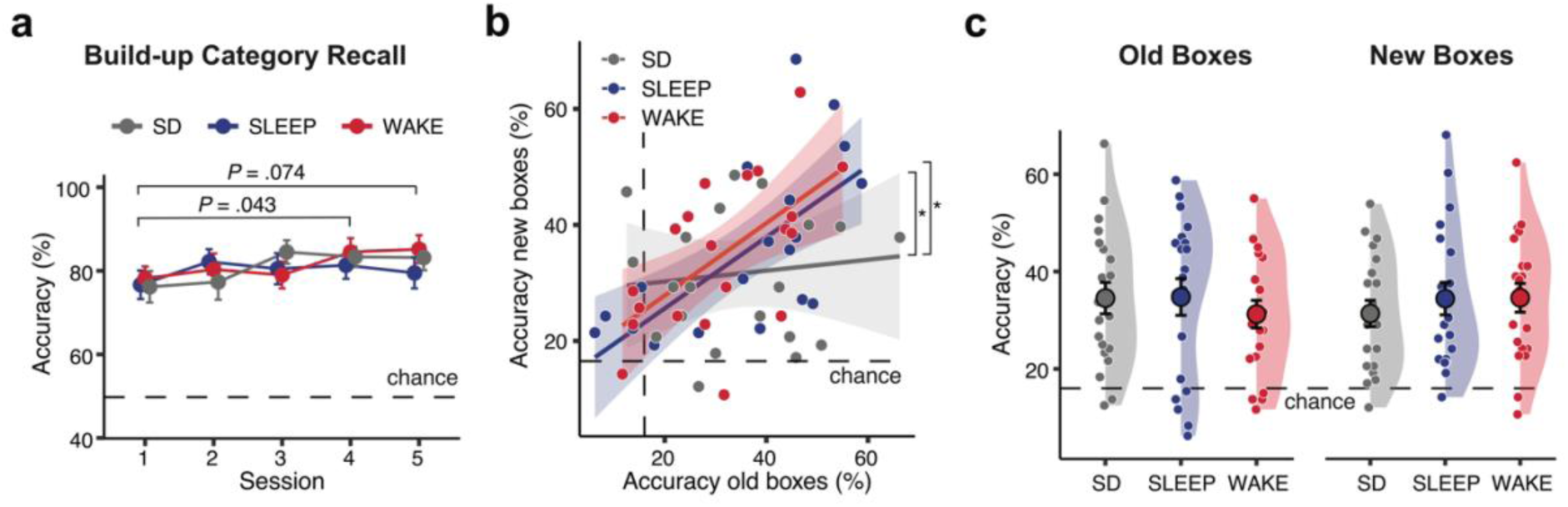
Encoding performance and the time-dependent relationship between episodic and schema memory. a, Category recall accuracy across the five schema build-up sessions. Performance was comparable across conditions (control, grey; sleep, blue; deprivation, red) and improved modestly in later sessions relative to session 1 (session 4: *P* = .043; session 5: *P* = .074). Dashed line, chance level (50%). Points indicate condition means ± SEM. **b,** Correlation between old-box and new-box recall accuracy indicating spatial interpolation as a prerequisite of schema use during retrieval. After the three-day delay, old box memory predicted interpolation performance in both the Sleep (*r* = 0.62, *P* < .001) and Wake (*r* = 0.66, *P* < .001) conditions, but not in the Short-delay condition tested on the same evening (*r* = 0.06, *P* = .709). Lines show linear regression fits with shaded 95% confidence bands. Dashed lines indicate chance performance. Asterisks indicate differences between regression lines at P < .05. **c,** Accuracy of category-ratio recall at schema retrieval for old boxes (left) and new boxes (right) locations. Accuracy was above chance level (16.67%, dashed line) and did not differ across conditions or old/new box locations. Violin plots show the distribution of individual participants; large circles with error bars denote condition means ± SEM. SD = Short-delay.

Central to our main hypothesis is the relationship between the location-object memories for the old boxes, and the spatial interpolation to the new box location at schema retrieval. If participants formed a spatial schema based on the learned category-ratio distribution, then better memory for old boxes should predict better interpolation to boxes at new locations, because both draw on the same underlying spatial representation. By contrast, if participants relied solely on item-level episodic memory without abstracting a spatial rule, performance for old and new box locations should be independent (**see Fig. 1c**). Indeed, memory for old boxes, measured by their associated spatial category ratio, predicted spatial interpolation performance for new boxes only after the three-day delay (**Fig. 2b**; Old Accuracy x Condition: *F*_1,53_ = 3.29, *P* = .045, η² = 0.11, follow-up Sleep vs. Short-Delay: *t_53_* = 2.93, *P* = .026; Wake vs. Short-delay: *t_53_* = 2.15, *P* = .037). In the Sleep condition, participants with better episodic recall for old box locations also showed better spatial interpolation (*r* = 0.62, *P* < .001); the same held for the Wake condition (*r* = 0.66, *P* < .001) condition. By contrast, participants in the Short-delay condition, tested in the same evening after schema build-up, showed no such relationship (*r* = 0.06, *P* = .709). This dissociation suggests that after a multi-day delay, participants used their episodic knowledge to infer category ratios at new locations; an important indicator for successful spatial schema formation.

In general, accuracy of category-ratio recall for old and new box locations did not differ between the retention conditions, old and new locations (Condition x Old/New: *F*_2,57_ = 1.05, *P* = .357; Old/New: *F*_1,57_ = 0.01, *P* = .917; Condition: *F*_2,56_ = 0.12, *P* = .886), or genders (*F*_1,56_ = 0.28, *P* = .597). Accuracies were above chance level in all conditions (16.67%; **Fig. 2c**; Sleep: *t*_19_ = 5.14; P < 0.001; Wake: *t*_19_ = 5.14; *P* < .001; Short-delay: *t*_19_ = 5.53; *P* < .001). This pattern indicates that there were no floor effects after the three-day delay relative to the Short-delay group.

### Sleep improves local spatial integration

To investigate the specific contribution of sleep to spatial interpolation performance, we computed a Schema Memory Index (SMI) that captures the global relational structure of the spatial distribution of object categories. The index represents a spatial measure of schema formation, which cannot be captured by simple correlations between average item-level accuracies that, although differentiating the Short-delay conditions from the two conditions with 3-day delays (see **Fig. 2b**), lack this structural information of the schema. The SMI was based on weighted graphs from the participants’ responses in which each box represented a node, with the edges connecting each new box to its surrounding old boxes and the edge weights defined by the difference in toy/household object counts between connected boxes (**Fig. 3a**). The SMI quantifies the structural similarity between the participants’ response graphs and the actual spatial distribution of item categories (see Methods for details).

**Fig. 3.**
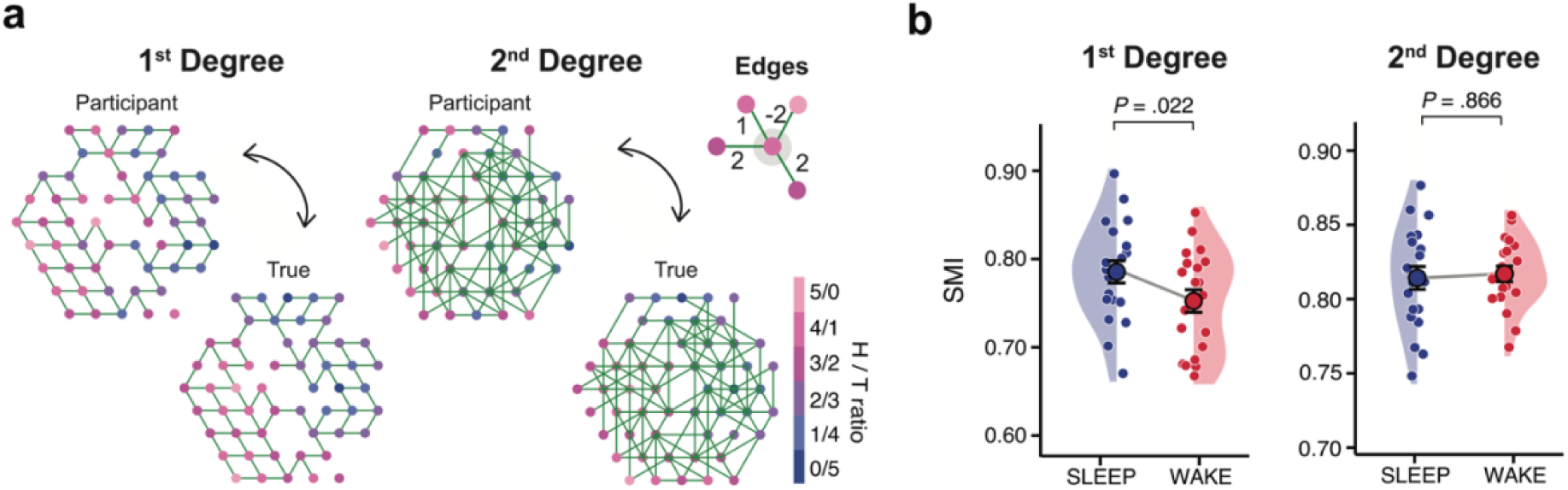
Sleep enhances local spatial integration. **a**, Graph-based quantification of schema memory. For each participant, weighted response graphs were constructed by connecting responses for new boxes to their neighbouring old box response, and weighting edges by the absolute difference in reported toy counts. Example participant vs. ground truth for 1^st^-degree (direct neighbors) and 2^nd^-degree (next-nearest neighbors) graphs. The node colour indicates the household-to-toy ratio. Inset (top right): illustration of edge computation for a single new box (pink) and its old-box neighbors, with edge weights reflecting the difference in toy counts between connected boxes. The global structure of participant and true graphs was compared using the normalized Laplacian spectral distance, from which the Schema Memory Index (SMI = 1 − spectral distance) was derived. **b**, SMI for the Sleep and Wake conditions. Sleep produced a higher SMI for 1^st^-degree graphs capturing the local spatial rule (*P* = .022), but not for 2^nd^-degree graphs (*P* = .866). Violin plots show individual participants; large dots with error bars denote condition means ± SEM.; grey lines connect condition means.

To test whether participants had acquired the local spatial rule (see **Fig. 1c**) underlying the spatial distribution, we calculated SMIs for 1^st^-degree graphs, capturing the local rule structure, and for 2^nd^-degree graphs as a control (**Fig. 3a**). Because the spatial rule was exclusively defined based on the direct old box neighbors, only SMIs for the 1^st^ degree graphs were expected to sensitively reflect effects on schema formation. The 2^nd^-degree graphs connected new boxes to their next-nearest old neighbors, which were visible from each new box location and thus available as potential cues regulating the participant’s responses. However, 2^nd^ degree neighbors were not used to define the rule underlying the pre-defined distribution. Hence, if sleep specifically enhanced learning of the local spatial rule, the sleep benefit should be evident for 1^st^-degree but not necessarily for the 2^nd^-degree graphs. We restricted these analyses to the Sleep and Wake conditions, as only these conditions showed significant spatial interpolation based on our correlational pre-analysis (**Fig 2b**).

Indeed, the SMI was higher in the Sleep than in the Wake condition for 1^st^-degree graphs capturing the local spatial rule (*t*_38_ = 2.38, *P* = .022, *d* = 0.75), but not for 2^nd^-degree graphs (*t*_38_ = 0.17, *P* = .866; **Fig. 3b**). This dissociation indicates that sleep specifically enhanced integration at the spatial scale at which the object category distribution rule was defined. In other words, participants who slept after encoding, were able to better reproduce the local relational structure of the category-ratio distribution, compared to participants who were sleep-deprived.

### Spatial schema is associated with shorter cue fixation

We investigated free-viewing eye movements during schema retrieval as an indicator of schema memory in an exploratory analysis. If participants had effectively formed a spatial representation of the category-ratio distribution, they may rely to lesser extent on active exploration of local cues and need less time to assess the objects relevant for navigation during retrieval but instead draw on their integrated spatial knowledge. We examined the relative proportion of fixations participants directed at task-relevant objects, that is, the ratio of gaze samples on relevant objects (response boxes and landmarks) to samples on irrelevant background (terrain and foliage). We also examined the Shannon entropy of the gaze trajectories. Both measures increase as a function of orienting and exploration and were, indeed, moderately correlated (*r* = 0.39, VIF = 1.18). Shorter object fixation durations predicted higher SMI (*r* = −.40, *P* = .012; **Fig. 4a**), while gaze entropy showed a trend-level association (*r* = −.30, *P* = .059; **Fig. 4b**). The relationship between fixation duration and schema memory remained significant when controlling for gaze entropy (*b* = .001, *P* = .049), with no difference between the Sleep and Wake conditions (*b* = .007, *P* = .413). Thus, participants with stronger schema memory spent less time inspecting local spatial cues during retrieval, consistent with increased reliance on internalized spatial representations as opposed to resampling the environment.

**Fig. 4.**
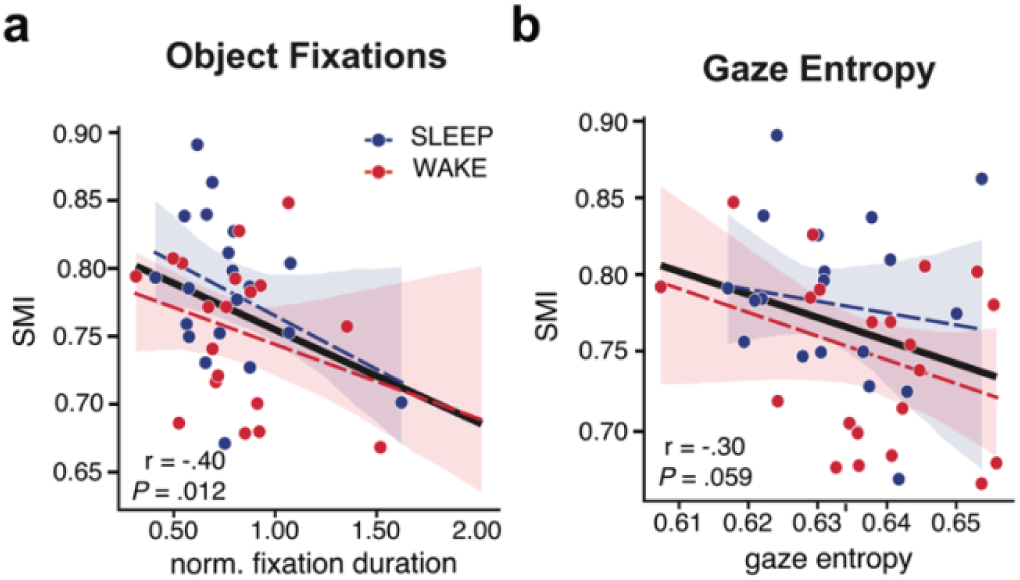
Eye-movement correlates of local spatial integration. **a**, Correlation between normalized object fixation duration and SMI across sleep (blue) and deprivation (red) conditions. Shorter fixation durations on objects during schema retrieval predicted higher SMI (*r* = −.40, *P* = .012). **b**, Correlation between gaze entropy and SMI. Lower gaze entropy showed a trend-level association with higher SMI (*r* = −.30, *P* = .059). The black line indicates the overall regression; coloured dashed lines show condition-specific fits with shaded 95% confidence bands.

### Slow oscillation-spindle coupling over frontal regions predicts schema memory

According to the active systems consolidation theory, the hippocampal-neocortical dialogue underlying the transformation of episodic memories into more generalized neocortical long-term representations is essentially orchestrated by SOs that, by nesting spindles in their excitable up-state, open windows of enhanced cortical plasticity to promote the formation of persisting neocortical representations ^32,45^. In this regard, the medial prefrontal cortex (mPFC) has been identified as a hub region for schema formation ^40^. We therefore tested whether SO-spindle coupling specifically over frontal cortical regions during the first post-encoding night predicted spatial schema memory as indexed by the SMI (for sleep macroarchitecture, see **Table S1**; for spindle and SO properties, see **Table S2**).

Peri-event time histograms of spindle occurrence rates locked to the SO negative half-wave peak at channel Fz confirmed the characteristic coupling pattern. Spindle occurrence was enhanced 400-800 ms after the SO trough, corresponding to the SO up-state (t_sum_ = 5.17, *P* < .001, *d* = 1.16; 1D cluster-corrected permutation test, 5,000 iterations; **Fig. 5a**). Time-frequency analysis of 12-16 Hz spindle power in this 400-800 ms window revealed that power was markedly higher during coupled SO events (those co-occurring with a spindle) than during non-coupled SOs occurring in isolation (*t_sum_ =* 4.49, *P* < .001, *d* = 0.63, **Fig. 5b**).

**Fig. 5.**
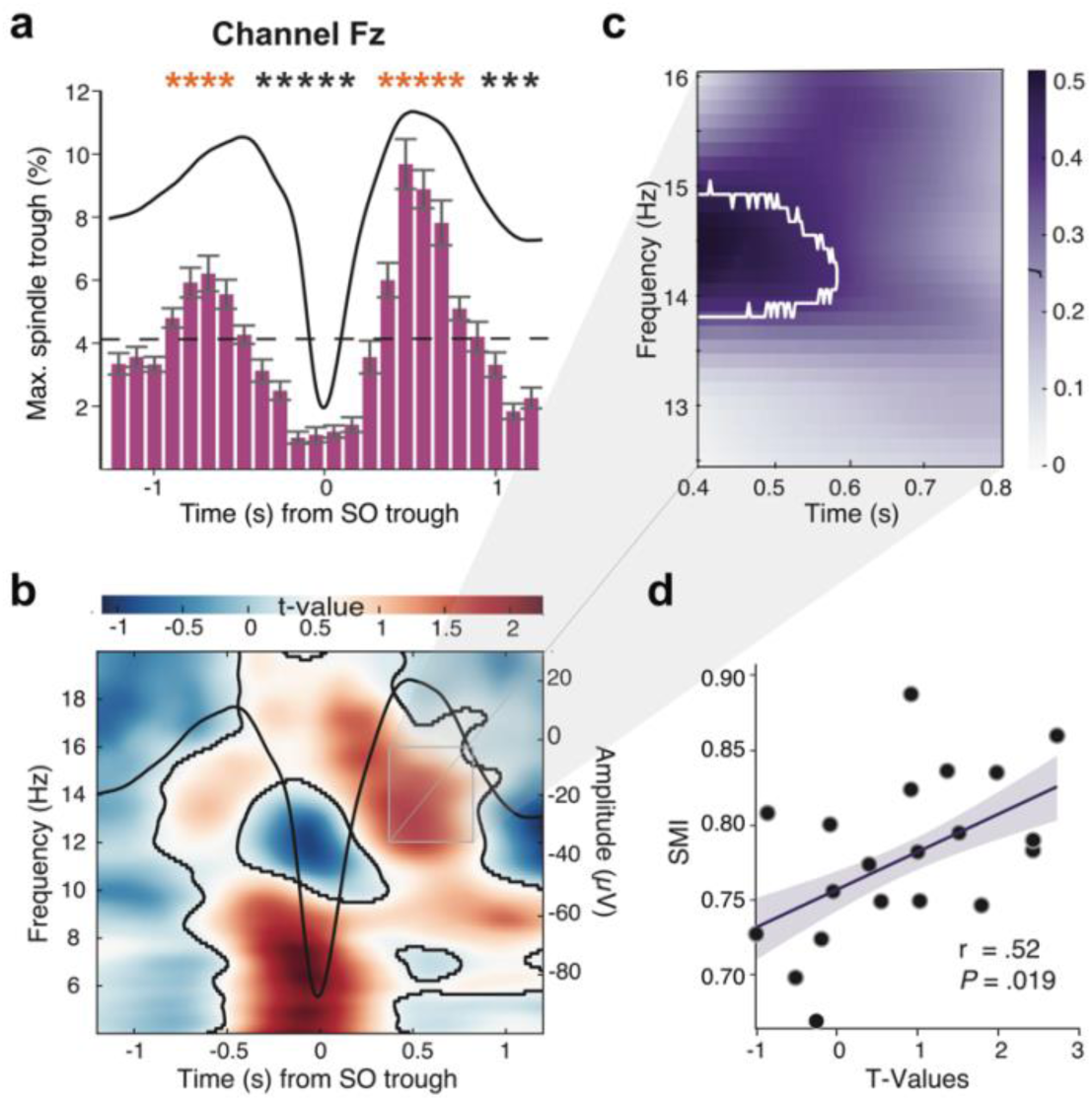
Frontal slow oscillation-spindle coupling during the first post-encoding night predicts schema memory. **a**, Peri-event time histogram of maximum spindle trough occurrence time-locked to the negative half-wave peak (0 s) of slow oscillations (SOs) at channel Fz. Black line shows average SO waveform. The characteristic coupling pattern is evident as enhanced spindle occurrence 400-800 ms after the SO trough, corresponding to the SO up-state. Red asterisks indicate time bins significantly exceeding the pre-event baseline (1D cluster-corrected permutation test, 5,000 iterations). Error bars denote SEMs; dashed line indicates the baseline spindle occurrence rate. **b**, Time-frequency representation of the *t*-statistic contrasting spindle-band power during SO events coupled versus not coupled with a spindle at Fz. Warm colours indicate greater power during coupled events. Black contour lines demarcate significant clusters (cluster-corrected permutation test). The SO waveform (black line, right axis) is overlaid for temporal reference. The grey rectangle marks the apriori time-frequency window of interest (400-800 ms, 12-16 Hz) used for the correlation with schema memory. **c**, Correlation map (Pearson *r*) between the coupled-minus-non-coupled power difference and the SMI within the 400-800 ms, 12-16 Hz window. The white contour delineates the significant cluster at 13.8-15 Hz (*r* = 0.42, *P* = .013). Darker shading indicates stronger positive correlations. **d**, Scatterplot of the maximum correlation between SO-spindle coupling power (*t*-values) and SMI at the peak time-frequency point (*r* = 0.52, *P* = .019). Line shows the linear regression fit with shaded 95% confidence band.

To test whether the strength of this coupling predicted schema memory, we correlated the power difference between coupled and non-coupled SO events with the SMI at each bin in the time-frequency spectrum (temporal range 400-800 ms, spectral range 12-16 Hz). We found a positive cluster spanning 13.8-15 Hz on the frequency axis and 400-600 ms after the SO negative half-wave peak (cluster *r* = 0.42, *P* = .013; **Fig. 5c**), with a maximum correlation of *r* = 0.52 (*P* = .019; **Fig. 5d**). Thus, participants who showed stronger frontal SO-spindle coupling during the first post-encoding night, subsequently also exhibited a higher SMI, linking time-locked sleep oscillatory coupling to spatial schema formation.

Control analyses confirmed the regional and temporal specificity of this effect. SO-spindle coupling over central (Cz; **Fig. S3a-d**) and parietal (Pz; **Fig. S3e-h**) channels in the Sleep condition did not correlate with the SMI (all cluster-corrected P > .468). To test whether this regional dissociation was reliable, we compared the coupling-SMI correlations across channels within the frontal cluster window (13.8-15 Hz, 0.4-0.6 s post SO-trough). The correlation was significantly stronger at Fz than at Pz (r_Fz_ = 0.42 vs. r_Pz_ = −0.16; Williams’s T2_15_ = 2.02, *P* = .03), with the same trend relative to Cz (r_Cz_ = −0.002; T2_15_ = 1.69, *P* = .056). Similarly, frontal SO-spindle coupling during the first recovery night in the Wake condition did not correlate with schema memory (**Fig. S3i-l**; cluster-corrected *P* > .432). Together, these findings support the selective relevance of the first post-encoding night for schema memory consolidation and highlight the specific role of frontal cortical oscillatory coupling in this mechanism.

## Discussion

This study shows that sleep, in addition to time, contributes to the de novo formation of a spatial schema. Participants learned a spatial distribution of object-category ratios across multiple learning sessions retrieved this information either after a short 30-min wake delay or longer 3-day delays. Within the 3-day delay, participants either spent the night after learning asleep or awake. We used spatial rule learning to measure successful schema memory formation. On a behavioral level, we found that participants used spatial interpolation to novel box locations only after a delay of three days and not at immediate recall. Importantly, spatial information integration after three days was better when sleep immediately followed schema build-up compared to a night of sleep deprivation. Electrophysiologically, this improved schema memory was associated with the precise coupling of spindles to the up-state of SOs after the build-up phase. This effect was spatially localized over frontal cortical regions. This evidence suggests that on top of passage-of time effects, sleep plays an active role in spatial schema memory formation.

Schema research in humans has mostly investigated how pre-existing schemas are shaped by new information through assimilation or accommodation^2,8,46^. This present study addresses the de novo formation of a spatial schema from episodic experience. This distinction is important because it shifts the focus from the top-down influence of prior knowledge on the learning of new information^47^ to the bottom-up process by which regularities across episodes are abstracted into a generalized spatial representation. Foundational rodent work on schema memory was conducted in the spatial domain^17,18^, yet research in humans has predominantly relied on non-spatial schemas^9,48,49^. Our VR paradigm bridges this gap by providing a controlled, large-scale spatial environment in which spatial rules can be embedded across multiple learning episodes without prior familiarity, directly connecting to spatial tasks used in animal models^7^.

A central finding of this study is that the passage of time was necessary for participants to form the schema, i.e., to interpolate from their spatial knowledge about old box locations to new locations. Specifically, schema-guided recall as indicated by the correlation between old-box and new-box recall performance, emerged only after a three-day delay, and not after the 30-min-delay retrieval test. This temporal dependence is consistent with theoretical accounts proposing that the transformation of episodic representations into schema-like knowledge is a gradual process^3,28,31^. Our data support this view: while recall of episodic memory information, i.e., the category-ratio for the individual old boxes, was comparable across all conditions, only after a multi-day delay was this episodic knowledge available for spatial generalization. This observation aligns with evidence that representations of regularities can require extended intervals to emerge even when sleep shortly after encoding initiates this process^50,21,51^. Graves and colleagues (2022) used iEEG to show that statistical regularities that were originally represented in the hippocampus, shifted to the mPFC over the course of three weeks.

Eye-tracking data provided an indirect measure in support of the acquired schema. Participants with better schema memory spent less time fixating on local spatial cues and tended to engage in less exploratory eye movements during retrieval. This pattern is consistent with schema-guided behavior: Once regularities are extracted, they can guide navigational decisions efficiently without requiring explicit attendance^8^.

Crucially, passage of time alone was not sufficient for optimal schema formation. Although both sleep and sleep-deprived participants showed a correlation between old-box and new-box performance as a marker of interpolation after the three-day delay, sleep specifically enhanced the integration of episodic spatial information, i.e., the category-ratio information associated with each old box into a coherent spatial schema. The selectivity of this effect is noteworthy: The sleep advantage emerged for 1^st^-degree graphs capturing the local spatial rule, but not for 2^nd^-degree graphs, indicating that sleep specifically facilitated integration at the spatial scale which was used to define the rule. Again, this benefit occurred without any improvement in recall of episodic spatial information (i.e., the category-ratio associated with the individual old boxes), arguing against a general consolidation account and in favor of sleep specifically facilitating the abstraction and integration process that defines schema formation^1,3,33,52^.

Overall, the present results converge with a growing body of evidence across species demonstrating that sleep actively transforms memory representations beyond simple stabilization^27,53,54^. In rats, only animals that slept during a 2 h post-encoding window exhibited schema memory for a spatial rule 24 h later, whereas animals kept awake did not^35^. According to active systems consolidation theory, the transformation of episodic into schema memory results from the repeated reactivation of episodic traces in hippocampal networks^28^, with the reactivation of overlapping episodic traces strengthening shared features, while weakening idiosyncratic details, fostering the abstraction of structures and the integration into schemas^27,34^ that are permanently stored in the neocortex^25,29^. Our findings extend this framework to spatial schema formation in humans: Sleep does not merely strengthen individual episodes but promotes the extraction of the spatial rule that connects them.

The finding that SO-spindle coupling over frontal cortical regions during the first post-encoding night predicted spatial schema memory provides a candidate mechanism linking sleep physiology to schema formation. A body of evidence in rodents and humans support the notion that the hippocampal-to-neocortical representational redistribution that underlies memory transformation during sleep, is orchestrated by the temporal coupling of neocortical slow oscillations (SOs), thalamocortical spindles, and hippocampal ripples during slow-wave sleep ^29,52^. Spindles nesting within SO up-states open windows of enhanced cortical plasticity thought to enable the strengthening of neocortical connections^55,56^. Our observation that this coupling specifically over frontal, but not central or parietal, cortex regions predicted schema memory is consistent with a substantial body of evidence implicating the medial prefrontal cortex (mPFC) as a hub for schema representation^2,8,15,35,51,57^.

In humans, retrieval of schema-congruent information engaged the mPFC and its connectivity with distributed cortical regions^49^. Most compellingly, intracranial recordings in humans demonstrated that hippocampal ripples during rest synchronize with mPFC activity, to predict the emergence of grid-like schema representations in entorhinal and prefrontal cortex^43^. Together with those findings, our results suggest that frontal SO-spindle coupling reflects the hippocampal-prefrontal dialogue through which episodic representations are transformed into integrated spatial schemas during sleep. To be noted, unlike SO-spindle coupling during sleep after the build-up phase of the Sleep group, this coupling did not predict schema memory when it occurred during the first recovery night in the sleep-deprived Wake group. This pattern reinforces the notion of a critical post-encoding time window for sleep-dependent schema consolidation, in line with evidence that the first sleep period after encoding is particularly important for memory consolidation and for the initiation of long-term abstraction processes^50,58,59^.

Given the conceptual importance of the hippocampal-prefrontal cortical dialogue underlying schema memory formation, it is a clear limitation of our study that it did not include direct measures of hippocampal activity. Although EEG-derived SO-spindle coupling likely reflects hippocampal-cortical communication^55^, the precise hippocampal contribution to schema formation in our task remains to be established. Another limitation pertains to the use of a virtual rather than a real-world environment. Although VR engages hippocampal and entorhinal spatial circuits, indicated by place-cell-like activity^19^ and grid-like representations ^60^, our task lacked real-world physical locomotion. Differences in hippocampal theta dynamics between virtual and real movement have been reported^61^ with real locomotion typically affording stronger multisensory integration and allocentric spatial encoding. Thus, our setup did not capture fully embodied spatial processing. However, it did allow head movements and body rotation. Also, video gaming experience and gender may have influenced spatial attention^62^, although we restricted participation to individuals with minimal video game experience (<1 session per month), and we did not see any gender effect. Importantly, if our data underestimate hippocampal engagement relative to real-world navigation, the sleep-dependent schema effects observed here could be even more pronounced in more naturalistic settings. Finally, stress following sleep-deprivation needs to be considered as potentially confounding factor in our Wake group. While we allowed for two recovery nights to exclude any stress-related effects on retrieval testing, future studies need to examine whether conditions of quiet wakefulness, in the absence of any stress, may invoke schema forming effects similar to those of sleep^63^.

In conclusion, this study provides converging evidence that the formation of spatial schema in humans is a time-dependent process that is actively supported by sleep. Further studies are indicated to clarify the underlying mechanisms, especially the role the hippocampus plays for abstracting spatial schemas during sleep.

## Materials & Methods

### Participants

60 healthy young adults (30 female; range 18-35 years, *mean* = 23.5 years) participated in the study. Participants were screened via a structured telephone interview. Inclusion criteria were normal visual acuity, normal color vision, and limited video-game experience (fewer than one gaming session per month on average). Exclusion criteria were current or past psychological, psychiatric or neurological diagnosis or treatment; medication use other than hormonal contraceptives, vitamin supplements, or thyroid medication; surgical procedures within the preceding three months; susceptibility to motion sickness; acute alcohol or drug influence; shift work or intercontinental travel (>4 h time-zone difference) within the preceding four weeks; irregular sleep-wake schedules (habitual sleep onset outside the 22:00-24:00 h window or habitual sleep duration outside 6-10 h); and participation in other sleep-manipulation studies within the preceding four weeks. The study was approved by the ethics committee of the University of Tübingen Medical Faculty. All participants provided written informed consent, including separate consent for eye-tracking recordings, and received financial compensation.

### Experimental procedure

We performed stratified block randomization to assign participants to one of three between-subjects conditions: sleep (Sleep, *N* = 20), sleep deprivation (Wake, *N* = 20) or immediate control (Short-delay, *N* = 20) with a fixed 50/50 male-to-female ratio within each group. Participants in the Sleep and Wake conditions completed an adaptation night in the sleep lab approximately one week before the first experimental session.

The experiment comprised three phases: schema build-up, a retention interval (sleep, sleep deprivation or 30-min break), and schema retrieval (see **Fig. 1d**).

#### Schema build-up

All conditions followed an identical encoding protocol. Participants arrived at the sleep lab at 16:30 h and completed entry questionnaires assessing current health status, stress, and sleepiness, followed by a baseline psychomotor vigilance test (PVT). They then donned the virtual-reality headset and completed a 15-min guided familiarization phase, in which they explored the hexagonal arena under experimenter guidance, practiced free navigation using the hand-held controller, and interacted with four box stimuli that would later serve as the test objects. The four boxes were placed in the center of the arena at locations irrelevant for the subsequent schema memory task.

Following a 5-min rest, participants began the schema build-up phase, which comprised five consecutive learning-retrieval cycles separated by 15-min breaks. Each cycle consisted of a learning block (∼30 min) and an immediate retrieval block (∼15 min). During learning, 34 boxes were distributed across fixed positions in the arena. On each trial, participants freely navigated to a box, which opened to reveal either a household item or a toy presented for 5 s. After presentation, participants categorized the object. Each learning session used a different start location within the arena to promote allocentric spatial encoding (see **Fig 1a**). During immediate retrieval, participants were sequentially spawned in front of each box, indicated the category of the object that had been presented at that box during the preceding learning session, and then selected the correct object from a pair of alternatives. Auditory feedback (correct/incorrect) was provided on each retrieval trial for both answers to reinforce accurate category-location and object-location associations. Between learning-retrieval cycles, participants completed questionnaires for 5 min (Stanford Sleepiness Scale, SSS; Multidimensional Mood Questionnaire, MDBF; Motion Sickness Assessment Questionnaire, MISC), performed a 5-min PVT and rested for 5 min. Schema build-up ended at approximately 20:40 h, followed by a standardized snack (two butter pretzels and an apple).

#### Retention interval

After schema build-up, participants entered one of three retention conditions. In the Sleep condition, polysomnographic electrodes were applied (∼60 min) and participants slept in the lab from 23:00 to 07:00 h. The following morning, participants completed sleep-quality questionnaires (see Questionnaires) and were fitted with a wrist-worn actigraph (see Actigraphy) to monitor activity and compliance with sleep-wake instructions over the subsequent days. They were discharged at 08:00 h and returned for schema retrieval three days after encoding, following two recovery nights at home during which they maintained their habitual sleep-wake rhythm and avoided napping during the day. A sleep diary was completed each morning during the inter-session interval, based on which participants were excluded if their sleep-wake pattern did not adhere to the instructions.

In the Wake condition, participants remained awake in the laboratory throughout the night under experimenter supervision. During the deprivation period (∼21:10-07:00 h), participants engaged in structured activities designed to maintain wakefulness without providing new spatial learning: board games, computer games (Snood, Tetris), audiobook listening, supervised walks and standardized snacks. After the wake night, participants completed questionnaires and were discharged at ∼08:00 h with an actigraph device. They were explicitly instructed not to nap during the following day. On the evening of the same day (18:00 h), they returned to the laboratory for a recovery night with polysomnographic recording (lights off at ∼19:30 h, lights on at 07:00 h), to control for non-specific effects of sleep on subsequent schema retrieval. Following an additional recovery night at home, participants returned for schema retrieval three days after encoding (at 14:30 h). Actigraphy data were reviewed upon return to verify compliance.

In the Short-delay condition, no electrodes were applied and no overnight stay was required. Following schema build-up and the standardized snack, participants rested for approximately 30 min before proceeding directly to schema retrieval on the same evening (from ∼21:15 h).

#### Schema retrieval session

After completing entry questionnaires, participants performed a single retrieval block (∼30 min). The arena now contained 54 boxes: the 34 original (old) boxes and 20 additional (new) boxes at previously unoccupied locations. Participants were spawned at the arena center and freely navigated to each box based on their choosing. For each box, they first responded whether it was a new or old box. For old boxes, they reported the category ratio (number of toys out of five objects) they had encountered during schema build-up. For new boxes, the instruction was: “infer the category ratio of the target box based on the box’s spatial location in the arena”, providing a measure of spatial interpolation, our indicator for successful schema memory formation. Following retrieval, participants responded to a last set of questionnaires assessing their VR experience, learning strategy, and awareness of the spatial rule. Finally, participants were then debriefed and the experiment was concluded.

### Virtual-reality setup

The VR system ran on a high-performance gaming PC equipped with an NVIDIA GeForce RTX 3090 GPU, an Intel Core i9-12900K processor, 32 GB DDR4 RAM (Corsair Vengeance LPX, 3200 MHz) and a Samsung 980 PRO 1 TB SSD. Participants navigated in a standing position using a hand-held Xbox controller and wore in-ear headphones for auditory feedback and immersive sounds, including random steps. While physical head and body rotations were possible, participants could move in the environment only using the controller. The virtual environment was developed in Unreal Engine 5.0 (Epic Games) and presented through a Varjo Aero headset (Varjo Technologies, Helsinki, Finland) running Varjo Base software v4.14 via SteamVR. The Varjo Aero headset provided integrated eye tracking with gaze data recorded at the display refresh rate. The target frame rate was maintained at approximately 80 frames per second to minimize motion sickness.

#### Hexagonal arena

The arena consisted of a large hexagonal outdoor space (100 m virtual diameter) that participants traversed with a maximal experienced speed of 8 km/h. The arena contained up to 54 box positions (34 during schema build-up and 54 during schema retrieval) arranged in a regular hexagonal grid. The hexagonal geometry ensured that each box had equidistant direct neighbors among varying spatial axial degrees, a property essential for unbiased spatial interpolation. The environment was enriched with distal spatial cues visible from all locations (green hills to the north, forest to the east, snowy mountains to the south and a village to the west) as well as a variety of proximal landmarks (houses, trees, a campground, a crop field, cars, wheelbarrows, a lake, a windmill, and a sawmill). These were positioned such that both direct neighbors and second-degree neighbors of any given box were visible from its location. The sky and all spatial cues remained constant across sessions. Footstep sounds were played during navigation to enhance immersion.

#### Stimuli

Object stimuli were three-dimensional models sampled from commercially available asset packages on the Unreal Engine Marketplace, representing either household items or toys (**Fig. 1b**). The 150 unique stimuli were validated in a pilot study (N = 10) using PsychoPy ^64^, in which participants classified images of each object. Only objects that were not incorrectly categorized by more than two participants were included in the main experiment. Each of the 34 old boxes contained five objects across schema build-up, drawn from the two categories with varying ratios (ranging from 5 toys/0 household items to 0 toys/5 household items), creating a continuous spatial gradient of category ratios across the arena (see **Fig. 1b**). The category ratio at each new-box location was defined by the rounded mean ratio of its directly adjacent old-box neighbors (see **Fig. 1c**). The objects presented at each box in each session during schema build-up were randomized across participants in each condition but matched between conditions.

#### Behavioral logging

Continuous frame-level logs recorded the following variables for each data frame: frame number, timestamp, player location and rotation (XYZ), velocity (XYZ), frames per second, head location and rotation (XYZ), gaze origin and direction (XYZ), and the identity of the fixated object (obtained via real-time ray tracing based on gaze position, see below for details). Trial-level logs captured all task-relevant events and responses. For learning trials, logged variables included trial and session number, trial timing, start location and rotation, movement onset, box identity, box arrival and opening times, presented object identity and category, object presentation duration, and category response. For immediate-retrieval trials, logged variables additionally included category-recall accuracy, object-recognition choice (left/right) and accuracy. For schema-retrieval trials, logged variables included box-type response (old/new) and accuracy, category-ratio response and accuracy, and all associated timestamps.

### Polysomnographic recordings

Polysomnographic recordings were obtained with a sampling rate of 500 Hz (BrainAmp, Brain Products, Gilching, Germany) during the first post-encoding night in the Sleep condition and the first recovery night in the Wake condition. Nine scalp electrodes were mounted at positions Fz, F3, F4, Cz, C3, C4, Pz, P3, P4 according to the international 10-20 EEG system. Additionally, horizontal and vertical eye movements (EOG), electromyogram (EMG) on the chin, and electrocardiogram (ECG) were recorded. The recordings started shortly after the light was turned off.

### Psychomotor vigilance test

Sustained attention was assessed using a 5-min computerized psychomotor vigilance task (PVT)^44^. At irregular intervals, a red counter appeared on a black screen and began incrementing. Participants pressed the spacebar as quickly as possible once the counter started. Reaction time was computed as the interval between counter onset and button press. The PVT was administered at baseline (before familiarization) and after each of the five schema build-up sessions.

### Questionnaires

Self-reported sleepiness was measured using the Stanford Sleepiness Scale (SSS) administered between each learning-retrieval cycle and at schema retrieval. Mood was assessed using the Multidimensional Mood Questionnaire (MDBF). Motion sickness was assessed using the Motion Sickness Assessment Questionnaire (MISC). A final questionnaire at debriefing assessed experienced immersiveness, self-reported learning strategy (intuition versus explicit memory recall), and awareness of the spatial distribution rule.

### Actigraphy

Participants in the Sleep and Wake conditions wore a wrist-worn actigraph device (MotionWatch 8, CamNtech, Cambridge, UK) on the non-dominant wrist from the morning after the first experimental session until schema retrieval three days later. Participants were instructed to press the event marker twice at each transition (lights off, wake-up, removal, reapplication). Actigraphy data were reviewed upon return to verify compliance with the instruction to maintain a regular sleep-wake schedule and to refrain from daytime napping.

### Data Processing

#### Schema Memory Index

To quantify schema memory, we constructed weighted graphs from both the true category ratio distribution and each participant’s responses. In each graph, the 20 new boxes were the schema-relevant nodes, that were connected by edges to their directly neighboring old boxes (1^st^ degree: equidistant direct neighbors only; 2^nd^ degree: next-nearest neighbors). Edge weights between nodes were defined as the absolute difference in toy counts between connected boxes, *w*_*ij*_ =∣ *k*_*i*_ − *k*_*j*_ ∣, where *k*_*i*_ and *k*_*j*_denote the toy count (range 0-5) at boxes *i* and *j*, respectively. To compare the global structure of the true and participant’s response graphs, we computed the normalized Laplacian spectral distance between both graphs. We represented the graphs as weighted adjacency matrices. For a weighted adjacency matrix *W*, the normalized Laplacian is defined as

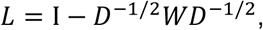

where *D* is the diagonal degree matrix, describing the number of edges to each node, and I the identity matrix. The spectral distance between two graphs with normalized Laplacian eigenvalues λ_1_, …, λ_*n*_ and 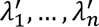 was computed as

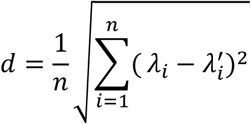

where *n* is the number of nodes. We used normalized eigenvalues to make the graph across neighborhood degrees (i.e. 1^st^ and 2^nd^ degree), with a different structure by construction, comparable. The Schema Memory Index (SMI) was then defined as

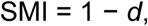

such that SMI = 1 reflects a perfect match between the response and true graphs and lower values indicate greater dissimilarity. This index was designed to capture whether participants had formed a coherent, integrated spatial representation of the category-ratio distributions, the hallmark of schema memory (**Fig. 6a-c**).

**Fig. 6.**
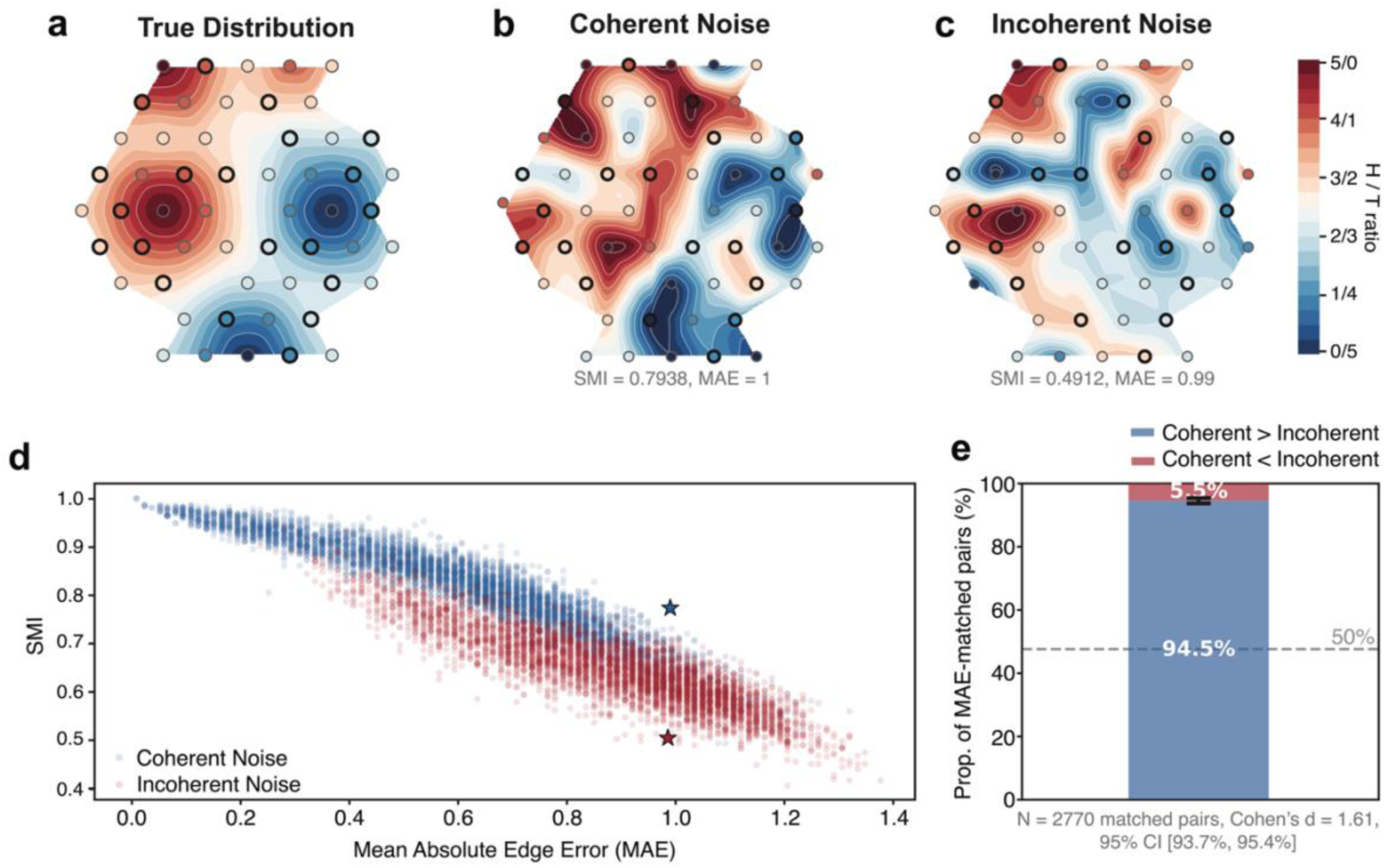
Simulation-based validation of the Schema Memory Index. **a**, Interpolated surface plot of the true category ratio distribution across the hexagonal arena. Circles represent individual box locations; colour indicates the number of toys (0-5) encountered across five learning sessions. **b**, Example simulated response with coherent noise, in which Gaussian noise was added to all box values, preserving the overall spatial gradient of the distribution. **c**, Example simulated response with incoherent noise, in which category ratios were randomly permuted among a subset of boxes, disrupting the spatial assignment while preserving the marginal distribution of ratio values. The coherent and incoherent examples were selected to have matched mean absolute edge error (MAE ≈ 1.0) but differ in SMI (0.79 versus 0.49). **d**, SMI plotted against MAE for 5,000 coherent (blue circles) and 5,000 incoherent (red circles) simulated responses. Stars mark the examples shown in **b** and **c**. At comparable MAE levels, coherent simulations tend to yield higher SMI values than incoherent simulations. **e**, Proportion of MAE-matched simulation pairs (tolerance < 0.1) in which the coherent response yielded a higher SMI than its incoherent counterpart (blue) versus the reverse (red). The dashed line indicates the 50% chance level. Error bar denotes the bootstrapped 95% confidence interval (10,000 iterations; *N* = 2,770 matched pairs).

To validate that the SMI captures the structural coherence of participants’ spatial representations beyond individual item-level accuracy, we performed simulations on two different types of response graphs: *Coherent noise* simulations added Gaussian noise (with a mean of zero and the standard deviations drawn uniformly between 0.3 and 2.5) to all box values simultaneously. This type of noise augmentation perturbed individual category ratios but preserved the overall spatial gradient of the distribution (**Fig. 6b**). *Incoherent noise* simulations randomly permuted category ratio values among a subset of boxes (30-100% of all boxes), which destroyed the spatial assignment of ratios to locations while preserving the marginal distribution of ratio values (**Fig. 6c**). For each incoherent simulation, a coherent simulation was generated with matched node-level error magnitude. We then paired coherent and incoherent simulations on mean absolute edge error (MAE; tolerance < 0.1), yielding 2,770 MAE-matched pairs from 5,000 simulations per class (**Fig. 6d**). If the SMI were insensitive to spatial structure and only reflected edge-level accuracy, the two classes should be indistinguishable at matched MAE. Instead, in 94.5% of matched pairs (**Fig. 6e**; 95% CI: 93.70-95.4%, bootstrapped over 10,000 iterations), the structurally coherent simulation yielded a higher SMI than its incoherent counterpart with a large effect size (Cohen’s *d* = 1.61), confirming that the SMI is sensitive to the global relational structure of category ratio distributions beyond what individual edge errors capture.

### Gaze Measures

#### Normalized cue fixation duration

We tracked which objects participants fixated during the experiment using ray tracing from the gaze position. This procedure gives per gaze sample the intersection of the gaze with objects in the VR arena. Each gaze sample was classified based on the name of the intersected object as either relevant (boxes and navigational landmarks) or irrelevant (background environment elements such as foliage and terrain). Samples with no reported intersection were excluded. Fixations on the trial-specific target box while participants were stationary were additionally excluded from all ratio computations, so that the measure reflects exploratory gaze behavior rather than the response duration for category ratios. We then computed, per trial, the ratio of gaze samples directed at task-relevant objects (chests and landmarks combined) to those directed at irrelevant background elements:

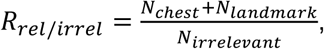

where *N_k_* denotes the number of frames classified as category *k*. Because both numerator and denominator scale linearly with trial duration, this ratio is inherently duration-invariant (confirmed empirically: Pearson *r* = 0.015 with trial duration). Higher values reflect proportionally greater allocation of time spent viewing navigation-relevant scene content.

#### Gaze transition entropy

To quantify the structure of exploratory scanning, we computed the Shannon entropy of the empirical distribution of object-to-object gaze transitions. Within each trial, consecutive frames fixating the same object were collapsed into a single viewing episode, and the resulting sequence of object changes was extracted. Each consecutive pair (*o_i_, o_i+1_*) constituted one transition, and the empirical probability of each unique ordered pair was estimated as its relative frequency:

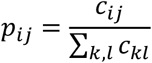

where *c_ij_* is the count of transitions from object *i* to object *j*. Shannon entropy over this distribution was computed as:

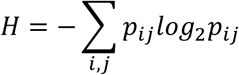

To account for differences in the number of unique objects fixated across trials, we normalized by the maximum possible entropy given n unique objects. Because self-transitions are structurally impossible in the transition sequence, the number of possible unique ordered pairs is *n*(*n* − 1), giving:

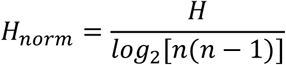

Values near 0 indicate highly structured scanning concentrated on a small number of object pairs; values near 1 indicate a near-uniform transition distribution, consistent with unsystematic exploration.

### Sleep scoring

Data were preprocessed in MATLAB 2024b (Mathworks Inc., Sherbom, Massachusetts) using the Sleeptrip toolbox (www.sleeptrip.org; RRID:SCR_017318). All EEG channels were re-referenced to averaged mastoids (M1 and M2). EEG and EOG data were band-pass filtered between 0.3 and 35 Hz, EMG between 10 and 100 Hz (4th-order Butterworth zero-phase filters). Sleep scoring was done visually by two experienced and independent scorers following standard criteria (AASM)^65^. All epochs on which the two raters disagreed were reviewed and matched between scorers. TST was calculated as time spent in stage 1, 2, SWS, and REM sleep. WASO was defined as the time participants were awake between sleep onset and final awakening (**Table S1**).

Channels were rejected based on visual inspection of the scoring epochs and the power spectra. In the Sleep condition, channel Cz was excluded for one participant (N = 19), and Pz for two participants (N = 18). For five participants in the Wake condition, channel Fz was excluded (N = 15). For the remaining channels, 30s-EEG epochs with artifacts labeled as movement arousals during sleep scoring were rejected from further analysis.

### Sleep event detection

#### Spindle detection

Spindles were detected using an automated algorithm implemented in Sleeptrip ^66^. Putative power peaks in the spindle frequency band (10-16 Hz) were visually detected from NREM epochs (i.e., N2 and N3). Before the inspection of the power spectra, the aperiodic 1/f^n^ component was removed from the signal by fitting a Lorentzian function using *specparam* ^67^. For each participant, spindle peaks were detected in the three midline channels (Fz, Cz, Pz; Sleep: 13.45 Hz ± 0.10; Wake: 13.34 Hz ± 0.09, mean ± SEM). This method was used to avoid the introduction of spurious spindle events ^68^. Since only half of the participants showed a detectable slow-spindle power peak over Fz, and all participants had a more prominent fast-spindle power peak over all three channels, we focused our analysis on the spindle frequency range of the more prominent faster peak. Thus, the signal of every channel was band-pass filtered in a ±1.5 Hz range around this peak (3 dB cutoff, 4th-order Butterworth zero-phase filter). The root-mean-square (RMS) representation of the signal was calculated at every sample point and smoothed using a moving average filter with a 200 ms sliding window. A potential spindle was marked if the amplitude of the smoothed RMS signal exceeded its mean by 1.5 SD of the filtered signal for 0.5-3 s. Local minima and maxima in the filtered spindle signal were marked as peaks and troughs. The largest trough was defined as the spindle peak time. The spindle amplitude was defined by the potential difference between the largest trough and the largest peak. Individual frequency of a concrete spindle was determined by adding the number of all troughs and peaks and dividing by twice the duration of the respective spindle. Spindles with boundaries closer than 0.25 s were merged.

#### Slow oscillation detection

Detection of slow oscillations at frontal electrodes was also based on an automated algorithm implemented in Sleeptrip and validated elsewhere ^30,69–71^. For all NREM epochs, the pre-processed EEG data was low-pass filtered at 4 Hz (6th-order Butterworth zero-phase filter). The whole signal was then divided into negative and positive half-waves that were separated by zero-crossings. A potential slow oscillation was defined as a negative half-wave followed by a positive half-wave in a frequency range between 0.5 and 1 Hz. Slow oscillation amplitude and frequency were computed as for spindles. Putative slow oscillations exceeding a trough of 1.25 times the mean trough of all putative slow oscillations as well as an amplitude of 1.25 times the average amplitude of all potential slow oscillations were accepted for further analysis.

#### SO-spindle coupling

To visualize co-occurrence of SOs and spindles, peri-event time histograms (PETHs) were constructed by counting spindle troughs in 100-ms bins over ±1.2 s around each SO trough, then normalizing by the total spindle count within that window. Statistical significance of the PETH structure was assessed via 1000 surrogate histograms generated by randomly shuffling bin assignments; surrogates were averaged per subject and channel to form a null distribution.

For event-locked cross-frequency coupling, we extracted artifact-free SO epochs (±3 s around each SO trough), band-pass filtered them at 0.3-1.25 Hz (6th-order Butterworth, zero-phase) and computed the instantaneous phase via the Hilbert transform. The same epochs were filtered in the average spindle band (12-16 Hz) and in 1Hz frequency bins of the spindle band, from which we derived the amplitude envelopes. To avoid edge artifacts, analyses were restricted to ±1.2 s around the SO trough. Within each event, channel, and subject, the maximal spindle amplitude (for the average spindle band and each frequency bin) and its corresponding SO phase was extracted.

Time-frequency dynamics of SO versus baseline epochs were examined using adaptive superlets^72^. The ±3 s epochs around SO-spindle events were paired with randomly selected, artifact-free baseline segments from the same sleep stage in which no spindle occurred during the SO. A fractional adaptive superlet transform was applied between 6 and 20 Hz in 0.25-Hz steps, using an initial Morlet base wavelet order of 3^rd^-order Morlet basewavelet with an order range from 5 to 15 multiplicatives, yielding enhanced time-frequency resolution compared to conventional wavelet methods.

### Statistical Analysis

All statistical analyses were performed in RStudio version 2023.12.1 using the *base* and *emmeans* packages, and MATLAB 2024b. The significance level was set to α = 0.05 using two-tailed testing unless specified otherwise. Post-hoc pairwise comparisons were corrected for multiple comparisons using the Holm method.

#### Schema build-up

Category recall accuracy was analyzed using a mixed analysis of variance (ANOVA) with the between-subjects factor Condition (Sleep, Wake, Short-delay) and within-subjects factors Session (1-5) and Object category (household, toy), with Gender as a covariate. Object recognition accuracy was analyzed with the same model structure. Self-reported sleepiness (SSS), mood (MDBF), motion sickness (MISC) and PVT reaction times were analyzed using two-way mixed ANOVAs with condition (Sleep, Wake, Short-delay) and session (baseline/sessions 1-5) as factors.

#### Schema retrieval

Category-ratio accuracy was analyzed using a mixed ANOVA with condition (Sleep, Wake, Short-delay) and box type (old, new) as factors and gender as a covariate. To test whether episodic memory predicted spatial interpolation, accuracy for new boxes was modelled as a function of mean accuracy for old boxes, condition and their interaction, with gender as a covariate. The SMI was compared between Sleep and Wake conditions for 1^st^-degree and 2^nd^-degree graphs using independent-samples *t*-tests. Pearson correlations were computed between the SMI and normalized cue fixation duration and gaze transition entropy. A multiple regression model tested the unique contribution of fixation duration while controlling for gaze entropy and condition.

#### SO-spindle coupling

For PETHs, we compared group-level bin counts against surrogate distributions using paired-sample t-tests at each time bin and channel. To correct for multiple comparisons, we employed cluster-based permutation testing (5,000 iterations).

Time-frequency representations (6-20 Hz) of SO and baseline trials over Fz, Cz and Pz in the Sleep condition, and over Fz in the Wake condition, were contrasted for each participant via independent-samples t-tests to produce t-maps of differences. To calculate the within-subject contrasts, we generated 100 random stimulus-baseline trial pairs (preserving the N2:N3 ratio). The t-maps were tested against zero on a group level using cluster-based permutation tests (5,000 permutations) in the ±1.2-s window around the SO trough. Finally, mean t-values within the spindle band (12-16 Hz) at 400-600 ms post-trough determined by the PETHs analysis were correlated with the SMI using Pearson’s *r*; and significance was determined via 2D-cluster-corrected bootstrap resampling (5,000 iterations).

To test the regional specificity of the association between frontal SO-spindle coupling and SMI, we compared the SMI-coupling correlation at Fz with those at Cz and Pz. For each participant, the mean coupled-versus-non-coupled spindle-band power difference was extracted from the time-frequency window of the significant Fz cluster (13.8-15 Hz, 0.4-0.6 s post SO-trough) at all three channels. Pairs of correlations (Fz vs. Cz; Fz vs. Pz) were then compared with Williams’s T_2_ test for dependent correlations sharing one variable ^73,74^. Tests were one-tailed, reflecting the directional a priori hypothesis that frontal coupling is more closely related to spatial schema formation than non-frontal coupling, consistent with the proposed role of medial prefrontal cortex as a hub for schema representation^2,8,43^.

## Data Availability

All main behavioral outcomes and preprocessed EEG datasets were deposited into the Open Science Framework (OSF) database and are available at the following URL: https://osf.io/8kg2j/.

## Code Availability

All custom computer code used to generate results that are central to the main claims in the manuscript are available on GitHub: https://github.com/LisaBastian/SchemAcS/.

## Acknowledgements

We thank Marcia Oliveira who supported data curation and Katharine A. Shapcott for advice on the VR-setup. This study was supported by grants from the Deutsche Forschungsgemeinschaft (DFG Bo 854/18-1) and the European Research Council to J.B. (ERC AdG 883098 Sleep Balance).

## Author Contributions (CRediT)

**L.B.:** Writing- original draft, Writing- review & editing, Visualization, Validation, Software, Methodology, Formal analysis, Data curation, Investigation, Conceptualization. **T.N.:** Writing-review & editing, Data curation, Software. **H.H.:** Data curation, Investigation. **K.R.:** Writing-review & editing, Project administration, Conceptualization. **J.B.:** Writing- review & editing, Validation, Resources, Project administration, Funding acquisition, Conceptualization.

## References

1. Bartlett, S. F. C. Remembering: A Study in Experimental and Social Psychology. (Cambridge University Press, 1995).

2. Gilboa, A. & Marlatte, H. Neurobiology of Schemas and Schema-Mediated Memory. Trends Cogn. Sci. 21, 618–631 (2017).

3. Ghosh, V. E. & Gilboa, A. What is a memory schema? A historical perspective on current neuroscience literature. Neuropsychologia 53, 104–114 (2014).

4. Farzanfar, D., Spiers, H. J., Moscovitch, M. & Rosenbaum, R. S. From cognitive maps to spatial schemas. Nat. Rev. Neurosci. 24, 63–79 (2023).

5. Moscovitch, M., Cabeza, R., Winocur, G. & Nadel, L. Episodic Memory and Beyond: The Hippocampus and Neocortex in Transformation. Annu. Rev. Psychol. 67, 105–134 (2016).

6. Robin, J. & Moscovitch, M. Details, gist and schema: hippocampal–neocortical interactions underlying recent and remote episodic and spatial memory. Curr. Opin. Behav. Sci. 17, 114–123 (2017).

7. Alonso, A., van der Meij, J., Tse, D. & Genzel, L. Naïve to expert: Considering the role of previous knowledge in memory. Brain Neurosci. Adv. 4, 2398212820948686 (2020).

8. Gilboa, A. & Moscovitch, M. No consolidation without representation: Correspondence between neural and psychological representations in recent and remote memory. Neuron 109, 2239–2255 (2021).

9. Van Buuren, M. et al. Initial Investigation of the Effects of an Experimentally Learned Schema on Spatial Associative Memory in Humans. J. Neurosci. 34, 16662–16670 (2014).

10. Sommer, T. The Emergence of Knowledge and How it Supports the Memory for Novel Related Information. Cereb. Cortex 27, 1906–1921 (2017).

11. Kim, M. & Maguire, E. A. Hippocampus, Retrosplenial and Parahippocampal Cortices Encode Multicompartment 3D Space in a Hierarchical Manner. Cereb. Cortex 28, 1898–1909 (2018).

12. Marchette, S. A., Vass, L. K., Ryan, J. & Epstein, R. A. Correction: Corrigendum: Anchoring the neural compass: coding of local spatial reference frames in human medial parietal lobe. Nat. Neurosci. 18, 926–926 (2015).

13. Zheng, L., Gao, Z., McAvan, A. S., Isham, E. A. & Ekstrom, A. D. Partially overlapping spatial environments trigger reinstatement in hippocampus and schema representations in prefrontal cortex. Nat. Commun. 12, 6231 (2021).

14. Tse, D. et al. Schemas and memory consolidation. Science 316, 76–82 (2007).

15. Tse, D. et al. Schema-dependent gene activation and memory encoding in neocortex. Science 333, 891–895 (2011).

16. Abdou, K. et al. Prefrontal coding of learned and inferred knowledge during REM and NREM sleep. Nat. Commun. 15, 4566 (2024).

17. Benchenane, K. et al. Coherent Theta Oscillations and Reorganization of Spike Timing in the Hippocampal- Prefrontal Network upon Learning. Neuron 66, 921–936 (2010).

18. Simon, K. C. et al. Sleep facilitates spatial memory but not navigation using the Minecraft Memory and Navigation task. Proc. Natl. Acad. Sci. U. S. A. 119, e2202394119 (2022).

19. Ekstrom, A. D. et al. Cellular networks underlying human spatial navigation. Nature 425, 184–188 (2003).

20. Betts, K. et al. An Examination of the Effects of Virtual Reality Training on Spatial Visualization and Transfer of Learning. Brain Sci. 13, 890 (2023).

21. Audrain, S. & McAndrews, M. P. Schemas provide a scaffold for neocortical integration of new memories over time. Nat. Commun. 13, 5795 (2022).

22. Sekeres, M. J., Winocur, G. & Moscovitch, M. The hippocampus and related neocortical structures in memory transformation. Neurosci. Lett. 680, 39–53 (2018).

23. Tompary, A. & Davachi, L. Integration of overlapping sequences emerges with consolidation through medial prefrontal cortex neural ensembles and hippocampal–cortical connectivity. eLife 13, e84359 (2024).

24. Winocur, G. & Moscovitch, M. Memory transformation and systems consolidation. J. Int. Neuropsychol. Soc. JINS 17, 766–780 (2011).

25. Diekelmann, S. & Born, J. The memory function of sleep. Nat. Rev. Neurosci. 11, 114–126 (2010).

26. Klinzing, J. G., Niethard, N. & Born, J. Mechanisms of systems memory consolidation during sleep. Nat. Neurosci. 22, 1598–1610 (2019).

27. Lewis, P. A. & Durrant, S. J. Overlapping memory replay during sleep builds cognitive schemata. Trends Cogn. Sci. 15, 343–351 (2011).

28. Lutz, N. D., Harkotte, M. & Born, J. Sleep’s contribution to memory formation. Physiol. Rev. 106, 363–483 (2026).

29. Staresina, B. P. et al. Hierarchical nesting of slow oscillations, spindles and ripples in the human hippocampus during sleep. Nat. Neurosci. 18, 1679–1686 (2015).

30. Bastian, L. et al. Spindle–slow oscillation coupling correlates with memory performance and connectivity changes in a hippocampal network after sleep. Hum. Brain Mapp. 43, 3923–3943 (2022).

31. Helfrich, R. F., Mander, B. A., Jagust, W. J., Knight, R. T. & Walker, M. P. Old Brains Come Uncoupled in Sleep: Slow Wave-Spindle Synchrony, Brain Atrophy, and Forgetting. Neuron 97, 221–230.e4 (2018).

32. Niethard, N., Ngo, H.-V. V., Ehrlich, I. & Born, J. Cortical circuit activity underlying sleep slow oscillations and spindles. Proc. Natl. Acad. Sci. 115, E9220–E9229 (2018).

33. Inostroza, M. & Born, J. Sleep for preserving and transforming episodic memory. Annu. Rev. Neurosci. 36, 79–102 (2013).

34. Stickgold, R. & Walker, M. P. Sleep-dependent memory triage: evolving generalization through selective processing. Nat. Neurosci. 16, 139–145 (2013).

35. Harkotte, M., Heimel, F., Dimitrov, S., Inostroza, M. & Born, J. The Emergence of New Schema Memory Requires Sleep. J. Neurosci. 46, e1810252025 (2026).

36. Feld, G. B., Bernard, M., Rawson, A. B. & Spiers, H. J. Sleep targets highly connected global and local nodes to aid consolidation of learned graph networks. Sci. Rep. 12, 15086 (2022).

37. Hennies, N., Lambon Ralph, M. A., Kempkes, M., Cousins, J. N. & Lewis, P. A. Sleep Spindle Density Predicts the Effect of Prior Knowledge on Memory Consolidation. J. Neurosci. Off. J. Soc. Neurosci. 36, 3799–3810 (2016).

38. Tamminen, J., Payne, J. D., Stickgold, R., Wamsley, E. J. & Gaskell, M. G. Sleep spindle activity is associated with the integration of new memories and existing knowledge. J. Neurosci. Off. J. Soc. Neurosci. 30, 14356–14360 (2010).

39. Contreras, M. P. et al. Context memory formed in medial prefrontal cortex during infancy enhances learning in adulthood. Nat. Commun. 15, 2475 (2024).

40. Cowan, E. et al. Sleep Spindles Promote the Restructuring of Memory Representations in Ventromedial Prefrontal Cortex through Enhanced Hippocampal-Cortical Functional Connectivity. J. Neurosci. Off. J. Soc. Neurosci. 40, 1909–1919 (2020).

41. Richards, B. A. et al. Patterns across multiple memories are identified over time. Nat. Neurosci. 17, 981–986 (2014).

42. Tang, W. & Jadhav, S. P. Sharp-wave ripples as a signature of hippocampal-prefrontal reactivation for memory during sleep and waking states. Neurobiol. Learn. Mem. 160, 11–20 (2019).

43. Xiao, Z. et al. Human hippocampal ripples align new experiences with a grid-like schema. Neuron 113, 3661–3672.e4 (2025).

44. Basner, M., Mollicone, D. & Dinges, D. F. Validity and Sensitivity of a Brief Psychomotor Vigilance Test (PVT-B) to Total and Partial Sleep Deprivation. Acta Astronaut. 69, 949–959 (2011).

45. Solano, A., Riquelme, L. A., Perez-Chada, D. & Della-Maggiore, V. Motor Learning Promotes the Coupling between Fast Spindles and Slow Oscillations Locally over the Contralateral Motor Network. Cereb. Cortex 32, 2493–2507 (2022).

46. Sherman, B. E. & Turk-Browne, N. B. Statistical prediction of the future impairs episodic encoding of the present. Proc. Natl. Acad. Sci. U. S. A. 117, 22760–22770 (2020).

47. van Kesteren, M., Ruiter, D., Fernández, G. & Henson, R. How schema and novelty augment memory formation. Trends Neurosci. 35, 211–9 (2012).

48. Brod, G., Lindenberger, U., Werkle-Bergner, M. & Shing, Y. L. Differences in the neural signature of remembering schema-congruent and schema-incongruent events. NeuroImage 117, 358–366 (2015).

49. van Kesteren, M. et al. Differential roles for medial prefrontal and medial temporal cortices in schema-dependent encoding: From congruent to incongruent. Neuropsychologia 51, 2352–2359 (2013).

50. Lutz, N. D., Diekelmann, S., Hinse-Stern, P., Born, J. & Rauss, K. Sleep Supports the Slow Abstraction of Gist from Visual Perceptual Memories. Sci. Rep. 7, 42950 (2017).

51. Graves, K. N. et al. Remembering the pattern: A longitudinal case study on statistical learning in spatial navigation and memory consolidation. Neuropsychologia 174, 108341 (2022).

52. Brodt, S., Inostroza, M., Niethard, N. & Born, J. Sleep-A brain-state serving systems memory consolidation. Neuron 111, 1050–1075 (2023).

53. Noack, H., Doeller, C. F. & Born, J. Sleep strengthens integration of spatial memory systems. Learn. Mem. Cold Spring Harb. N 28, 162–170 (2021).

54. Vorster, A. P. & Born, J. Sleep and memory in mammals, birds and invertebrates. Neurosci. Biobehav. Rev. 50, 103–119 (2015).

55. Helfrich, R. F. et al. Bidirectional prefrontal-hippocampal dynamics organize information transfer during sleep in humans. Nat. Commun. 10, 3572 (2019).

56. Latchoumane, C.-F. V., Ngo, H.-V. V., Born, J. & Shin, H.-S. Thalamic Spindles Promote Memory Formation during Sleep through Triple Phase-Locking of Cortical, Thalamic, and Hippocampal Rhythms. Neuron 95, 424–435.e6 (2017).

57. Hebscher, M. & Gilboa, A. A boost of confidence: The role of the ventromedial prefrontal cortex in memory, decision-making, and schemas. Neuropsychologia 90, 46–58 (2016).

58. Talamini, L. M., Nieuwenhuis, I. L. C., Takashima, A. & Jensen, O. Sleep directly following learning benefits consolidation of spatial associative memory. Learn. Mem. 15, 233–237 (2008).

59. Gais, S., Lucas, B. & Born, J. Sleep after learning aids memory recall. Learn. Mem. 13, 259–262 (2006).

60. Doeller, C. F., Barry, C. & Burgess, N. Evidence for grid cells in a human memory network. Nature 463, 657–661 (2010).

61. Garg, T. et al. The relationship between object-based spatial ability and virtual navigation performance. PLOS ONE 19, e0298116 (2024).

62. Feng, J., Spence, I. & Pratt, J. Playing an action video game reduces gender differences in spatial cognition. Psychol. Sci. 18, 850–855 (2007).

63. Wamsley, E. J. Memory Consolidation during Waking Rest. Trends Cogn. Sci. 23, 171–173 (2019).

64. Peirce, J. et al. PsychoPy2: Experiments in behavior made easy. Behav. Res. Methods 51, 195–203 (2019).

65. Berry, R. B. et al. AASM Scoring Manual Updates for 2017 (Version 2.4). J. Clin. Sleep Med. JCSM Off. Publ. Am. Acad. Sleep Med. 13, 665–666 (2017).

66. Weber, F. SleepTrip as a Tool to Create Analysis Pipelines for Investigations of Polysomnography in Clinical Populations. Int. J. Psychophysiol. 168, S65 (2021).

67. Donoghue, T. et al. Parameterizing neural power spectra into periodic and aperiodic components. Nat. Neurosci. 23, 1655–1665 (2020).

68. Ujma, P. P. et al. A comparison of two sleep spindle detection methods based on all night averages: individually adjusted vs. fixed frequencies. Front. Hum. Neurosci. 9, (2015).

69. Ngo, H.-V. V., Martinetz, T., Born, J. & Mölle, M. Auditory Closed-Loop Stimulation of the Sleep Slow Oscillation Enhances Memory. Neuron 78, 545–553 (2013).

70. Bastian, L. et al. Long-term memory formation for voices during sleep in three-month-old infants. Neurobiol. Learn. Mem. 215, 107987 (2024).

71. Kurz, E.-M., Bastian, L., Mölle, M., Born, J. & Friedrich, M. Development of slow oscillation–spindle coupling from infancy to toddlerhood. SLEEP Adv. 5, zpae084 (2024).

72. Moca, V. V., Bârzan, H., Nagy-Dăbâcan, A. & Mureșan, R. C. Time-frequency super-resolution with superlets. Nat. Commun. 12, 337 (2021).

73. Steiger, J. H. Tests for comparing elements of a correlation matrix. Psychol. Bull. 87, 245–251 (1980).

74. Williams, E. J. The Comparison of Regression Variables. J. R. Stat. Soc. Ser. B Stat. Methodol. 21, 396–399 (1959).

