## Supplementary Information for "Sleep enhances spatial schema memory formation in humans"

This PDF file includes:
Supplementary Results
Supplementary Figure 1 to 3
Supplementary Table 1 to 2
Link to Supplementary Video 1 to 3

**Supplementary Results**

Self-reported sleepiness during schema build-up was lower in the Wake condition, compared to two other conditions and, as expected, increased across learning sessions (**Fig. S2a**, Condition: $F_{2,57}$ = 3.31, $P$ = .044, η² = 0.10; Session: $F_{4,228}$ = 23.64, $P$ < .001, η² = 0.29). However, objectively measured sustained attention as assessed with the psychomotor vigilance test (Basner et al., 2011) did not differ between the retention conditions across encoding sessions and schema retrieval (**Fig. S2b**, Condition x Session: $F_{12,398}$ = 0.43, $P$ = .950, η² < 0.01). Although self-reported mood appeared to fluctuate differently across conditions and sessions (**Fig. S2c**, Condition x Session: $F_{8,228}$ = 2.38, $P$ = .018, η² = 0.07), no pairwise comparison reached significance. Motion sickness decreased across sessions (**Fig. S2b,** Session effect: $F_{4,228}$ = 4.40, $P$ = .002, η² = 0.07). Males and females experienced the environment similarly realistic across conditions (3.26/5 ± 0.23, $F_{2,49}$ = 0.48, $P$ = .622, η² = 0.02), and operating the gaming controller during the task was rated as highly intuitive (4.46/5 ± 0.16, $F_{2,49}$ = 0.01, $P$ = .991, η² = 0.29). Taken together, encoding performance prior to the sleep manipulations appeared to be well matched across conditions.

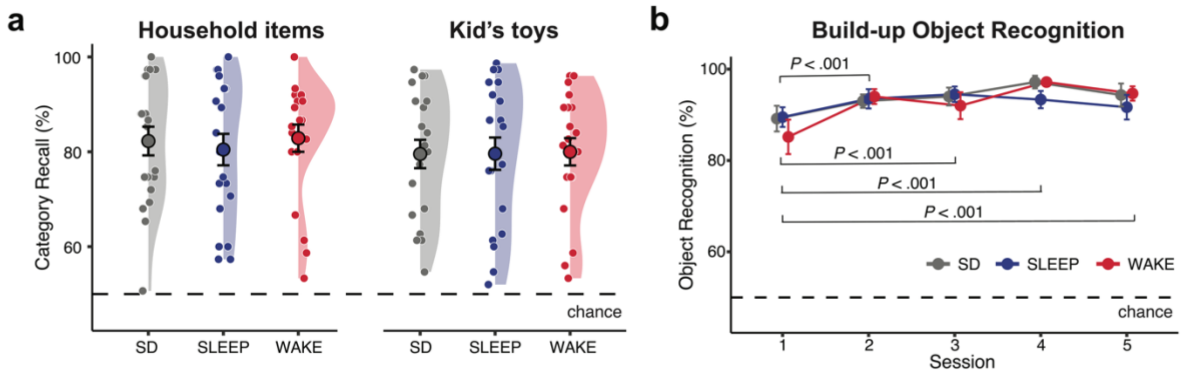


**Supplementary Fig. S1 | Schema build-up performance by object category and session. a**, Category recall accuracy during schema build-up, separated by object category (household items, left; kids' toys, right) and condition (SD, grey; SLEEP, blue; WAKE, red). Performance was comparable across conditions and object categories. Violin plots show individual participants; large circles with error bars denote condition means ± s.e.m. Dashed line indicates chance level (50%). **b**, Object recognition accuracy across the five schema build-up sessions. Participants performed worse in session 1 compared with all subsequent sessions (all *P* < .001), with no differences between conditions. Points show condition means ± s.e.m. Dashed line indicates chance level (50%).SD = Short-delay.

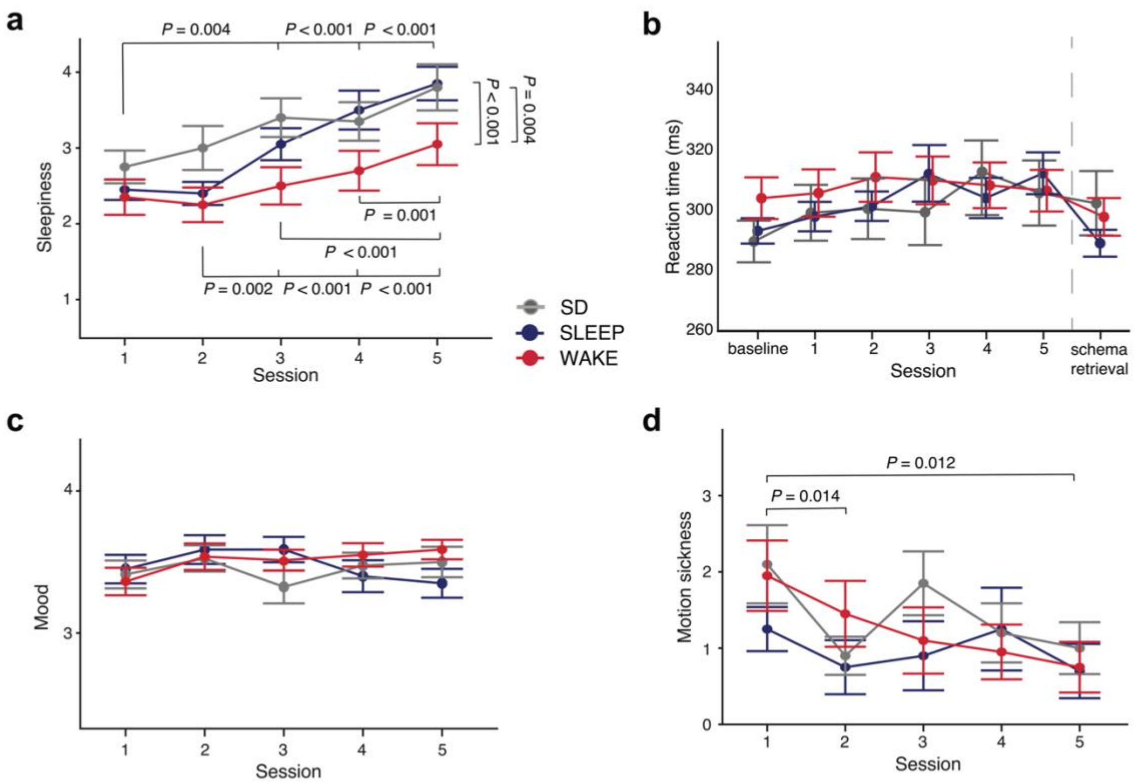


**Supplementary Fig. S2 | Control measures across schema build-up sessions. a**, Self-reported sleepiness (Stanford Sleepiness Scale) across sessions 1–5 for each condition (SD, grey; SLEEP, blue; WAKE, red). Sleepiness was lower in the deprivation condition and increased across sessions in all conditions. **b**, Self-reported mood across sessions. Although the Condition × Session interaction reached significance, no pairwise comparisons survived correction for multiple comparisons. **c**, Self-reported motion sickness across sessions. Motion sickness decreased across sessions. **d**, Psychomotor vigilance task (PVT) reaction times at baseline, across sessions 1–5 and before schema retrieval. Sustained attention did not differ between conditions. In all panels, points show condition means ± s.e.m. Significant pairwise comparisons (Holm-corrected) are indicated. SD = Short-delay.

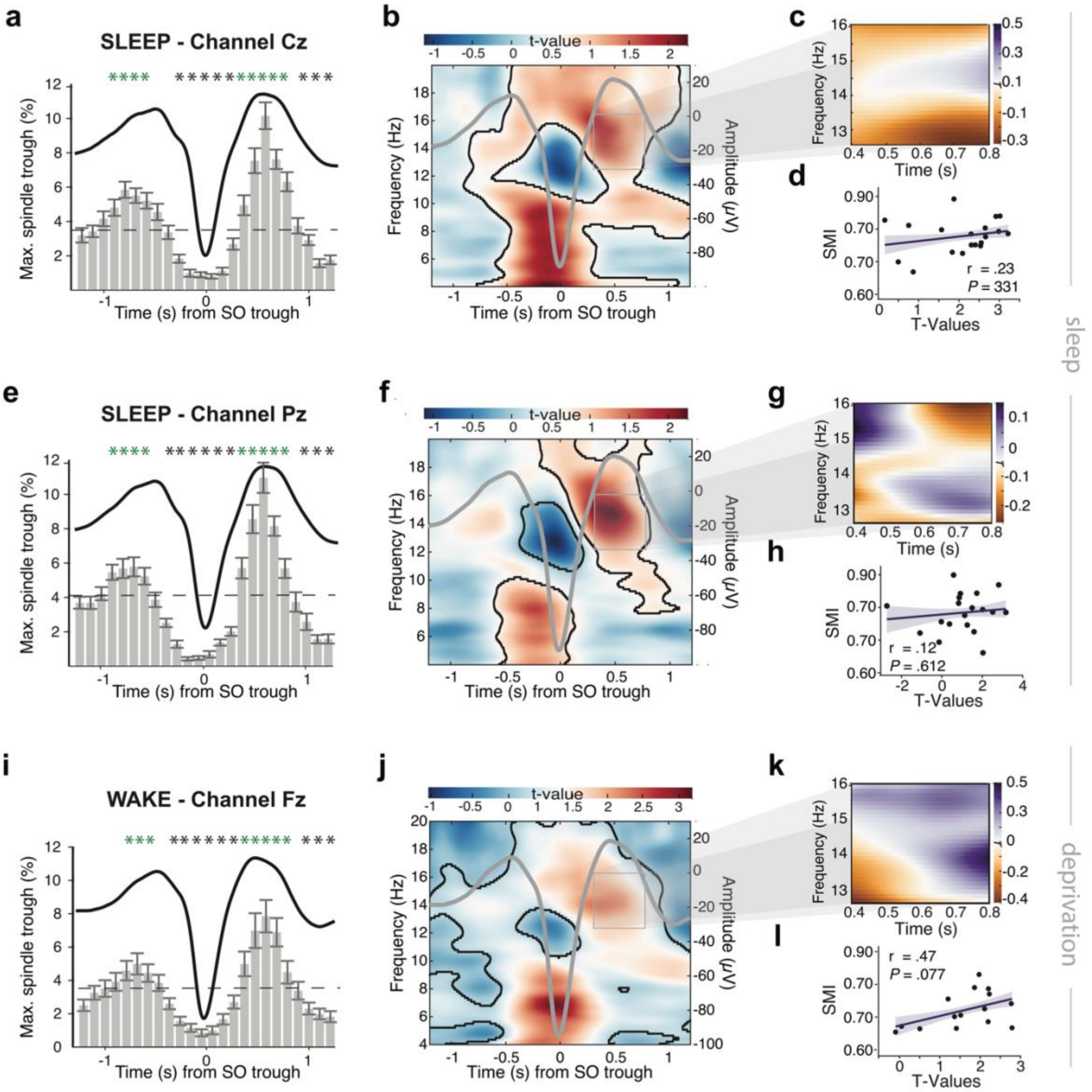


**Supplementary Fig. S3 | Control analyses for spindle–SO coupling over central and parietal regions and for the deprivation condition.** **a**–**d**, Sleep condition, channel Cz. **a**, Peri-event time histogram of maximal spindle trough occurrence rate time-locked to the SO negative half-wave peak. Green asterisks mark significant time bins (1D cluster-corrected permutation test). Error bars denote SEM. **b**, Time–frequency representation of *t*-statistics contrasting coupled versus non-coupled SO events. Black contours demarcate significant clusters; the SO waveform is overlaid (grey line, right axis). **c**, Correlation map (Pearson *r*) between the coupled-minus-non-coupled power difference and the SMI within the 400–800 ms, 13–16 Hz window. **d**, Scatterplot of SMI against peak coupling *t*-values ($r$ = .23, $P$ = .331). **e**–**h**, Sleep condition, channel Pz. Same layout as **a**–**d** ($r$ = .12, $P$ = .612). **i**–**l**, Wake condition, channel Fz (first recovery night). Same layout as **a**–**d** ($r$ = .47, $P$ = .077). Neither central nor parietal coupling in the sleep condition, nor frontal coupling during the first recovery night in the deprivation condition, was significantly associated with schema memory, supporting the selective role of frontal spindle–SO coupling during the first post-encoding night.

*Table S1.* Sleep macroarchitecture for the sleep and deprivation conditions during the respective first night of sleep following schema build-up (*N* = 20 per group)

| *Condition* | *Parameter* | *Mean (SD)* | *range* |
|---|---|---|---|
| Sleep | TST (min) | 483.70 (15.74) | 433.00 – 511.00 |
| | WASO (min) | 41.98 (27.16) | 19.50 – 135.50 |
| | N1 (min) | 22.63 (5.89) | 13.00 – 32.00 |
| | N2 (min) | 241.90 (29.46) | 181.50 – 305.50 |
| | N3 (min) | 80.35 (37.75) | 19.50 – 170.00 |
| | REM (min) | 96.85 (26.30) | 37.00 – 154.50 |
| | WASO (% of TST) | 8.73 (5.66) | 3.91 – 27.49 |
| | N1 (% of TST) | 4.67 (1.18) | 2.68 – 6.57 |
| | N2 (% of TST) | 49.98 (5.66) | 39.33– 62.22 |
| | N3 (% of TST) | 16.61 (7.71) | 3.97– 34.07 |
| | REM (% of TST) | 20.00 (5.44) | 8.55 – 31.89 |
| Wake | TST (min) | 654 (74.86) | 424.00 – 743.50 |
| | WASO (min) | 61.06 (71.97) | 5.00 – 268.50 |
| | N1 (min) | 2.37 (1.84) | 0.50 – 43.00 |
| | N2 (min) | 360.67 (83.16) | 73.00 – 456.50 |
| | N3 (min) | 103.94 (37.46) | 22.00 – 147.50 |
| | REM (min) | 113.31 (35.42) | 31.00 – 172.00 |
| | WASO (% of TST) | 10.24 (14.71) | 0.88 – 63.33 |
| | N1 (% of TST) | 1.42 (1.53) | 0.08 – 6.96 |
| | N2 (% of TST) | 54.49 (11.12) | 17.22 – 67.48 |
| | N3 (% of TST) | 15.85 (5.76) | 4.69 – 24.98 |
| | REM (% of TST) | 17.04 (4.47) | 7.31 – 23.19 |

*Note.* TST = total sleep time, WASO = wake after sleep onset, REM = rapid eye movement

*Table S2.* Spindle and SO characteristics for the sleep and deprivation conditions during the respective first night of sleep following schema build-up (*N* = 20 per group)

| Condition | Parameter | SOs | | | Spindles | | |
|---|---|---|---|---|---|---|---|
| | | *Fz* | *Cz* | *Pz* | *Fz* | *Cz* | *Pz* |
| Sleep | Mean frequency (Hz) | 0.86 (0.02) | 0.86 (0.02) | 0.84 (0.02) | 12.73 (0.41) | 13.03 (0.42) | 13.13 (0.45) |
| | Down-to-up slope (μV/s) | 768.71 (255.10) | 729.41 (194.58) | 624.29 (152.75) | --- | --- | --- |
| | Density (#/min) | 4.63 (1.11) | 4.27 (1.06) | 4.28 (1.10) | 4.90 (0.63) | 5.17 (0.63) | 5.61 (0.64) |
| | Amplitude (μV) | 162.66 (52.66) | 155.24 (42.27) | 136.63 (35.07) | 27.28 (7.90) | 31.62 (8.29) | 30.74 (7.48) |
| | Duration (s) | 1.16 (0.03) | 1.17 (0.02) | 1.18 (0.02) | 0.80 (0.03) | 0.83 (0.04) | 0.86 (0.05) |
| Wake | Mean frequency (Hz) | 0.88 (0.02) | 0.88 (0.02) | 0.85 (0.03) | 12.62 (0.40) | 13.01 (0.26) | 13.05 (0.36) |
| | Down-to-up slope (μV/s) | 870.15 (171.15) | 839.33 (141.46) | 720.85 (112.63) | --- | --- | --- |
| | Density (#/min) | 4.46 (1.19) | 3.91 (0.89) | 3.86 (0.87) | 4.92 (0.36) | 5.44 (0.38) | 5.71 (0.64) |
| | Amplitude (μV) | 175.21 (35.43) | 170.44 (25.88) | 149.74 (21.79) | 28.89 (6.23) | 37.27 (7.64) | 37.79 (8.83) |
| | Duration (s) | 1.12 (0.03) | 1.15 (0.03) | 1.18 (0.04) | 0.83 (0.04) | 0.85 (0.04) | 0.91 (0.06) |

*Note.* Values expressed as mean (SD).

**Supplementary Videos**

Video material showing example trials of a random participants during the learning session, the immediate retrieval session and the schema retrieval session can be found in the data repository at Open Science Framework (OSF; https://osf.io/8kg2j/files/osfstorage).
